# Powdery mildew infection induces a non-canonical route to storage lipid formation at the expense of host thylakoid lipids to fuel its spore production

**DOI:** 10.1101/2023.12.15.571944

**Authors:** J. Jaenisch, H. Xue, J. Schläpfer, E.R. McGarrigle, K. Louie, T.R. Northen, M.C. Wildermuth

**Author notes:** **Corresponding author’s email address**: Mary. C. Wildermuth. Authors contributed equally to this work (J. Jaenisch, H. Xue). Department of Plant and Microbial Biology, University of Zurich, Zurich 8008 (J. Schläpfer). The author(s) responsible for distribution of materials integral to the findings presented in this article in accordance with the policy described in the Instructions for Authors (https://academic.oup.com/plcell/pages/General-Instructions) is (are): Mary. C. Wildermuth.

## Abstract

Powdery mildews are obligate biotrophic fungi that manipulate plant metabolism to supply lipids, particularly during fungal asexual reproduction when fungal lipid demand is extensive. The mechanism for host response to fungal lipid demand has not been resolved. We found storage lipids, triacylglycerols (TAGs), increase by 3.5-fold in powdery mildew-infected tissue. In addition, lipid bodies, not observable in uninfected mature leaves, are present in both cytosol and chloroplasts at the infection site. This is concurrent with decreased thylakoid membrane lipids and thylakoid disassembly. Together, these findings indicate that the powdery mildew induces localized thylakoid membrane degradation to promote storage lipid formation. Genetic analyses show the canonical ER pathway for TAG synthesis does not support powdery mildew spore production. Instead, Arabidopsis DIACYLGLYCEROL ACYLTRANSFERASE 3 (DGAT3), shown to be chloroplast-localized and to be largely responsible for powdery mildew-induced chloroplast TAGs, promotes fungal asexual reproduction. Powdery mildew-induced leaf TAGs are enriched in thylakoid associated fatty acids, which are also present in the produced spores. This research provides new insights on obligate biotrophy and plant lipid metabolism plasticity and function. Furthermore, by understanding how photosynthetically active leaves can be converted into TAG producers, more sustainable and environmentally benign plant oil production could be facilitated.

## INTRODUCTION

As obligate biotrophic pathogens, powdery mildews acquire nutrients supplied by living host cells to support their life cycle and have specialized strategies for maximizing the output of these tissues (Glawe 2008; Wildermuth et al. 2017). In the *Arabidopsis thaliana-Golovinomyces orontii* interaction, the establishment of the fungal feeding structure, called a haustorium, occurs by 24 hours post inoculation (hpi). By 5 days post inoculation (dpi), asexual reproductive structures called conidiophores form. These conidiophores contain chains of conidia which store energy in the form of lipids and glycogen (Both et al. 2005; Micali et al. 2008). Thus, the fungal demand for nutrients is especially high during asexual reproduction. As a response to the nutritional demands, a metabolic switch occurs in the host infected leaves. Mature leaves are considered source tissues producing hexoses for transport to growing parts of the plant. However, powdery mildew infection induces localized signatures of mobilization of carbohydrates to the tissue underlying the fungal infection site, for fungal acquisition (Clark and Hall 1998; Sutton, Henry, and Hall 1999; Fotopoulos et al. 2003; Swarbrick, Schulze-Lefert, and Scholes 2006). Furthermore, localized transcriptome profiling using laser microdissection shows the expression of genes associated with enhanced glycolysis and respiration to be increased, while the expression of chlorophyll biosynthesis genes is decreased at the powdery mildew infection site, in support of a localized source to sink transition (Chandran et al. 2010). Analysis of powdery mildew genomes found reduced carbohydrate metabolism pathways but relatively complete fatty acid (FA) metabolism and utilization pathways, suggesting lipids may be a preferred nutrient (Liang et al. 2018). And, an early study found enhanced lipid accumulation in powdery mildew infected cucumber leaves compared to uninfected leaves (Abood and Lösel 1989).

Microbial acquisition of host lipids has emerged as a common strategy across host-microbe systems, particularly for obligate biotrophs including human intracellular pathogens (Atella et al. 2009; Costa et al. 2018). For plant obligate biotrophs, the arbuscular mycorrhizal fungi (AMF) symbiosis in which AMF colonize plant roots, providing minerals to the plant host while acquiring host sugars and lipids is best studied (MacLean, Bravo, and Harrison 2017; Luginbuehl et al. 2017; Kameoka and Gutjahr 2022). AMF induce a specific shift in host lipid metabolism, catalyzed by enzymes specific to plants colonized by AMF, to yield 2-monoacylglycerols (2-MG), with C16:0 2-MGs preferred. While 2-MGs appear to be the likely final product transferred to AMF, this has not been verified, and it is possible other lipids may also be transported particularly if acquisition is facilitated by exocytotic vesicles. Once these host lipids are acquired, they are remodeled by the AMF and stored primarily as TAGs in lipid bodies for future use.

By contrast with the AMF symbiosis, almost nothing is known about how the powdery mildew fungus manipulates host metabolism for fungal lipid acquisition. Because powdery mildews have the capacity to synthesize FAs, unlike AMF which are FA auxotrophs (Kameoka and Gutjahr 2022), we focus our studies on powdery mildew-infected leaves during powdery mildew asexual reproduction (5+ dpi), when spores replete with lipid bodies (Both et al., 2005) are formed. We reason that host lipids would be most in demand at this phase of the powdery mildew life cycle. Furthermore, Jiang and colleagues, as part of their research on AMF, show host FA manipulation is reflected in powdery mildew spore FAs (Jiang et al. 2017). Specifically, their introduction of UcFatB, a fatty acid thioesterase that terminates FA elongation early, terminating with C12:0, into Arabidopsis resulted in increased C12:0 FAs in both host leaves and powdery mildew spores.

Plant lipid metabolism is dynamic across developmental stages and responsive to environmental stimuli, modifying energy content of storage tissues, altering membrane fluidity at different temperatures, minimizing lipotoxicity, and providing chemical signals (Baud et al. 2008; Moellering, Muthan, and Benning 2010; Okazaki and Saito 2014; Cavaco, Matos, and Figueiredo 2021). In plants such as Arabidopsis, acyl-chains are produced in chloroplasts, with the exception of a small fraction generated in mitochondria, and their subsequent assembly into lipids occurs via pathways operating in the chloroplast (prokaryotic pathway) and the endoplasmic reticulum (eukaryotic pathway). In Arabidopsis leaves, approximately 38% of newly synthesized FAs are utilized in the prokaryotic lipid-synthesis pathway, whereas the remaining 62% are directed towards the eukaryotic pathway (Browse et al. 1986). A portion of acyl-chains from ER-assembled lipids are subsequently transported – likely as PA and/or DAG – back to the plastid to serve as substrates for thylakoid lipid synthesis (Yao et al. 2023; Hölzl and Dörmann 2019). Triacylglycerols (TAGs), neutral storage lipids with three fatty acids attached to a glycerol backbone, are packaged into lipid bodies. Eukaryotes synthesize TAGs in the ER via two major pathways: the Kennedy pathway and the acyl-CoA independent pathway (C. Xu, Fan, and Shanklin 2020). Diacylglycerol acyltransferases (DGAT, EC 2.3.1.20) catalyze the final and rate-limiting step in TAG synthesis forming TAG from diacylglycerol (DAG) and acyl-CoA in the Kennedy pathway. Whereas, phospholipid:diacylglycerol acyltransferase (PDAT, EC 2.3.1.158) catalyzes the final and rate-limiting step in acyl-CoA independent TAG synthesis with TAG formed from DAG and a phospholipid (PL) acyl donor, i.e. phosphatidylcholine (PC) remodeled from the Lands Cycle (Dahlqvist et al. 2000; Zhang et al. 2009; L. Wang et al. 2012).

In this study, we show powdery mildew-induced TAG accumulation in mature *Arabidopsis thaliana* leaves occurs at the infection site concurrent with powdery mildew asexual reproduction. We employ genetic, microscopic, and lipidomic approaches to uncover a non-canonical route for plant TAG synthesis via AtDGAT3 to support powdery mildew spore formation. AtDGAT3 is unusual in that, unlike the ER membrane proteins DGAT1 and DGAT2, it is a soluble metalloprotein containing a [2Fe-2S] cluster (Aymé et al. 2014). We show AtDGAT3 is localized to the chloroplast and responsible for plastidic TAG synthesis that occurs at the expense of thylakoid membranes. We further speculate on controls over functional roles of ER-versus chloroplast-derived lipid bodies in the powdery mildew interaction, powdery mildew acquisition of the chloroplast-derived lipid bodies, and controls over AtDGAT3 stability and activity. Our findings open further avenues of investigation with respect to biotroph-host interactions and plant response to stress or aging (e.g. leaf senescence). Moreover, this work could facilitate more sustainable production of vegetable oil, biofuels and other specialty chemicals (X.-Y. Xu et al. 2018).

## RESULTS

### Powdery mildew infection increases triacylglycerols in the host leaf while phospholipids decrease

Powdery mildew fungi are obligate biotrophs that rely entirely on the host for nutrients. Powdery mildew asexual reproduction creates a high metabolic demand for lipids as the powdery mildew feeding structures, haustoria, and newly formed spores are filled with lipid bodies at 5 dpi when asexual reproduction is first apparent **(Fig. 1, A-C**).

**Figure 1.**
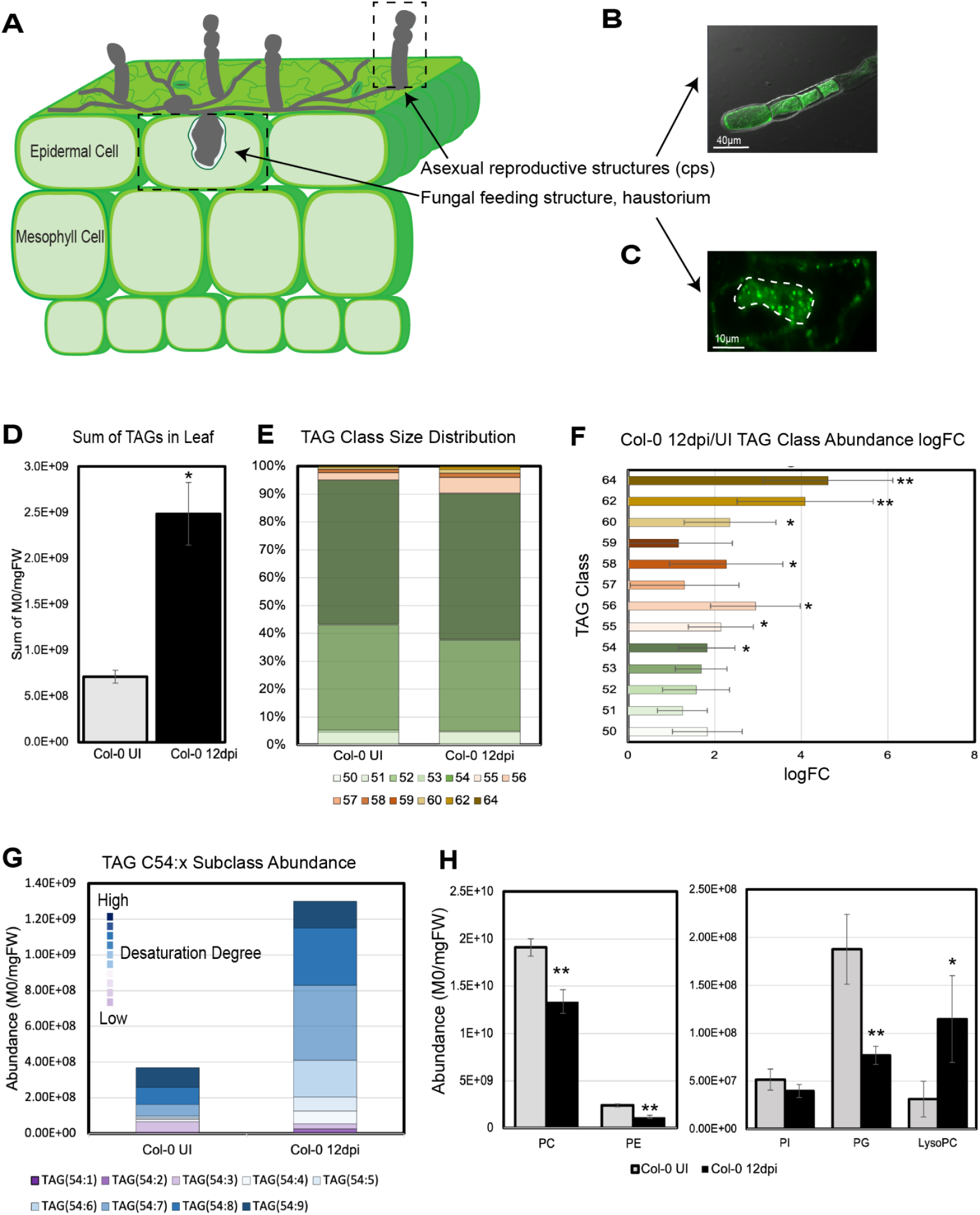
TAG abundance is increased in infected Col-0 leaves. A) Cross-section depicting powdery mildew infection of Arabidopsis leaf at 5dpi. B-C) BODIPY 505/515 neutral lipid-stained powdery mildew structures: B) Asexual reproductive structure, conidiophore (cp), bar= 40µm. C) haustorium, bar= 10µm, white dashed line outlines haustorium. D) Total TAGs (C50-C64) detected in uninfected (UI) and 12dpi leaf lipid extracts ±STD, n= 3. E) Distribution of TAG classes in UI and 12dpi leaf lipid extracts. F) Log_2_ fold change (LogFC) of TAG abundance by class in 12 dpi vs UI leaf lipid extracts ±STD, n= 3. G) Abundance of C54:x subclasses in UI and 12 dpi leaf lipid extracts. Assumes TAGs within this *m/z* range have similar desorption/ionization properties. H) Summed abundance of detected phospholipids (M0/mgFW) in UI (grey) and 12dpi (black) leaf lipid extracts ±STD, n= 3. Significance between UI and 12 dpi tested by 2-tailed T-test * p ≤ 0.05, ** p ≤ 0.01.

To understand how host lipid metabolism is manipulated to meet this fungal lipid demand, we performed lipid profiling of uninfected and parallel powdery mildew-infected leaves at 12 days post inoculation (dpi). This later time point exhibits sufficient powdery mildew proliferation to allow us to assess the impact of the powdery mildew in whole leaf analyses. Lipids were extracted and identified by LC-MS/MS fragmentation patterns (**Supplemental Fig. 1, Supplemental Dataset 1**). Our results show that TAGs increase in 12 dpi washed leaf extracts relative to uninfected leaf extracts, with a 3.5-fold increase in abundance (**Fig. 1D**). Overall, there is a shift to TAGs containing longer acyl chains, including very long chain fatty acids (VLCFA, ≥C20), assessed at ≥C56:x, which increase 7-fold with infection (**Fig. 1E-F, Supplemental Fig. S1; Supplemental Dataset 1**). Examination of the most abundant TAG class, C54:x, shows an increase of 3.5-fold with infection accompanied by a shift towards a more desaturated profile in infected leaves (**Fig. 1F-G**); this reflects increased 18:3 and 18:2 FA composition (**Supplemental Dataset 1**).

While TAGs increase, phospholipids decrease in abundance in extracts from infected leaves at 12 dpi compared to parallel uninfected leaves (**Fig. 1H, Supplemental Dataset 1**). Phosphatidylcholine (PC), the dominant phospholipid in mature Arabidopsis leaves, decreases by 30%. Phosphatidylethanolamine (PE) and phosphatidylglycerol (PG) decrease by 50% and 60% respectively in infected leaves. Although a net decrease in total phosphatidylinositol (PI) with infection of 20% is observed, it is not statistically significant. Lysophosphatidylcholines (LPC) increase by ∼4-fold in infected leaves. The observed decrease in PC is consistent with increased TAG synthesis utilizing DAG formed from PC (and PA) via DGATs. The decreases in the other phospholipids (PE, PI, PG) may facilitate increased flux to TAG accumulation. Furthermore, the indication that LPC increases at 12 dpi suggests possible operation of the Lands Cycle using PDAT1.

In summary, our data indicates that the powdery mildew remodels host lipid metabolism to promote localized TAG accumulation.

### Genetic analyses indicate the canonical route for plant TAG synthesis in the ER hinders powdery mildew asexual reproduction while chloroplast-localized DGAT3 promotes it

We next examined the impact of genes encoding proteins catalyzing the final and rate-limiting step in canonical TAG biosynthesis in the ER (Vanhercke et al. 2019) on powdery mildew spore production, *AtDGAT1 (At2g19450)*, *AtDGAT2 (At3g51520)*, and *AtPDAT1* (At5g13640), using Arabidopsis null mutants and/or spray-induced gene silencing (SIGS). Our employed SIGS methodology specifically silences targeted genes with minimal off targets (Methods; McRae et al. 2023). Furthermore, the Arabidopsis DGATs evolved independently, contain distinct functional domains, and share little sequence similarity (Yin et al. 2022). In seed oil accumulation, AtDGAT1 and AtPDAT1 play dominant roles. A null mutant in *AtDGAT1* shows a 30% reduction in seed TAGs, while RNAi silencing of *PDAT1* in a *dgat1-1* background or *DGAT1* in *pdat1-1* background results in 70 to 80% decreases in seed oil content (Katavic et al. 1995; Zhang et al. 2009). While it doesn’t contribute to seed TAG accumulation, AtDGAT2, along with AtDGAT1 and AtPDAT1, can impact leaf TAG accumulation (Zhou et al. 2013; Fan, Yan, and Xu 2013). Furthermore, we explored the impact of ATP-binding cassette A 9 (ABCA9), demonstrated to import FA/acyl-CoA into the ER and to exhibit a 35% reduction in seed TAG accumulation in the *abca9-1* mutant (Kim et al. 2013). The ER-localized long-chain acyl-CoA synthetase 1 (LACS1) was also investigated because it acts on long chain and very long chain FAs (Lü et al. 2009) which we observed to increase with infection (**Fig. 1E-F**) and is the only ER-localized LACS (Zhao et al. 2010) with enhanced expression at the powdery mildew infection site at 5 dpi (Chandran et al. 2010).

To our surprise, *dgat1-1* and *abca9-1* null mutants allow for enhanced powdery mildew spore production, 24% and 50% more, respectively, than wild type (WT) plants, whereas, the *lacs1-1* and *pdat1-2* mutants show no significant change in spore production (**Fig. 2A**). Knockdown of *AtDGAT1* via SIGS results in more than 60% increase in spore production, whereas, silencing of *AtDGAT2* shows no difference in spore production from mock treatment (**Fig. 2B**). Taken together, our findings indicate that TAG synthesis in the ER, using the FA importer ABCA9 and DGAT1, is not used to support powdery mildew spore production, but instead hinders it.

**Figure 2.**
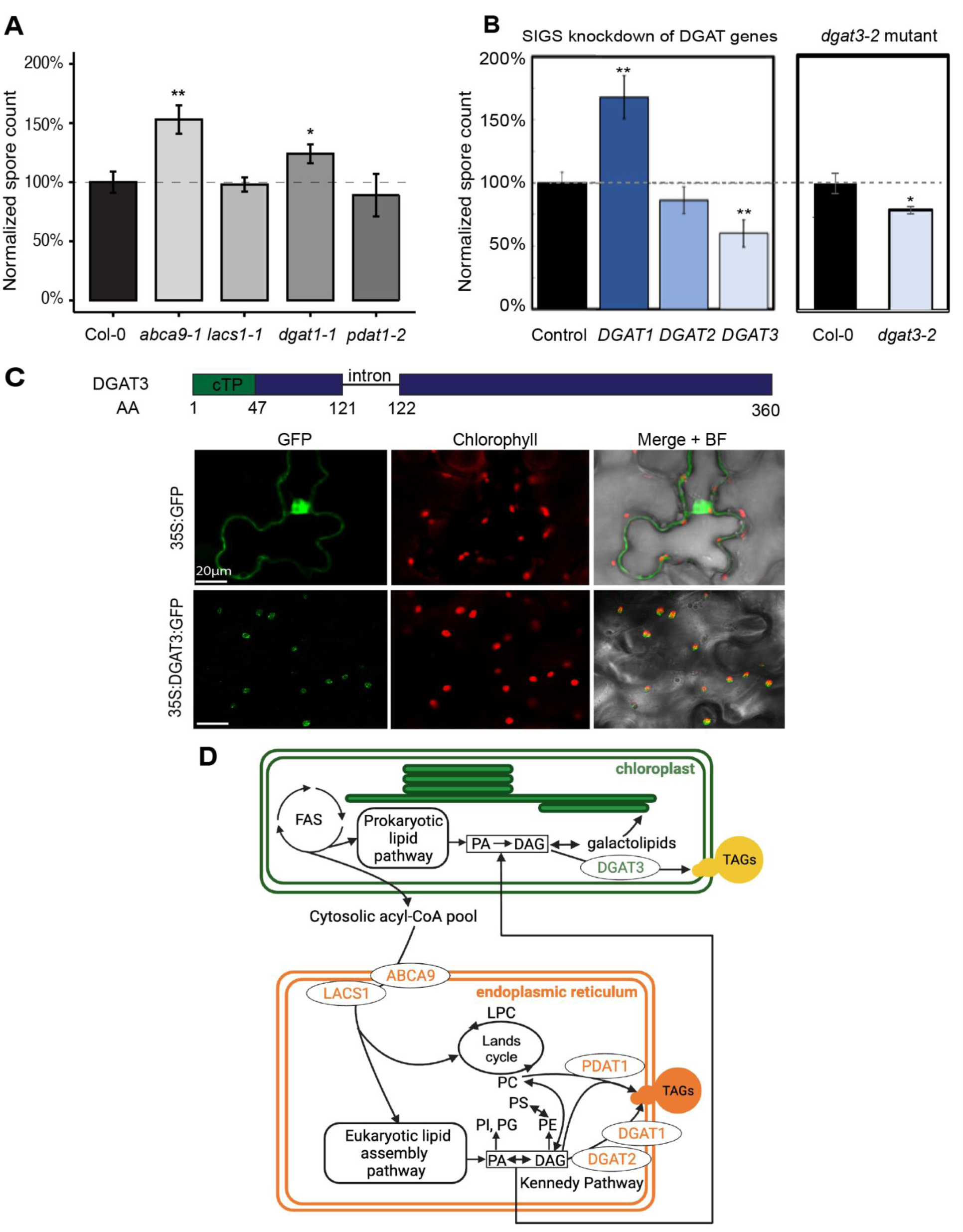
Canonical TAG synthesis in the ER hinders powdery mildew asexual reproduction while chloroplast-localized DGAT3 promotes it. A) Spore counts/mg leaf FW at 9dpi of mutants normalized to WT Col-0 for mutants involved in the canonical route for TAG synthesis in the ER (±STD, n= 5-8). B) Comparison of spore counts/mg leaf FW on WT plants with *DGAT* genes silenced via spray-induced gene silencing (SIGS) and *dgat3-2* mutant vs. WT at 9dpi (±STD, n= 4-8). Significance by 2-tailed T-Test *p ≤ 0.05, **p ≤ 0.01. C) AtDGAT3 protein is predicted to have a chloroplast transit peptide by the DeepLoc 2.0 and LOCALIZER program. Confocal microscopy images of transient expression of 35S:AtDGAT3-GFP in *Nicotiana benthamiana*. D) Simplified model of tested players that may have contributed to Arabidopsis TAG production. Abbreviations: ABCA, ATP-binding cassette A; BF, bright field; DAG, diacylglycerol; DGAT, Diacylglyceroltransferase; FAS, fatty acid synthase complex; LACS, long chain acyl-CoA synthetase; LPC, lysophosphatidylcholine; phosphatidic acid; PC, phosphatidylcholine; PE, phosphatidylethanolamine; PI, phosphatidylinositol; PS, phosphatidylserine; PDAT, phospholipid:diacylglycerol acyltransferase; TAGs, triacylglycerols. See Figure 1 for data on phospholipids.

While AtDGAT1 and AtDGAT2 are ER-localized and membrane-bound, the third Arabidopsis DGAT protein, AtDGAT3, contains a predicted N-terminal chloroplast transit peptide (cTP) and no transmembrane domain (Aymé et al. 2018). AtDGAT3 was initially shown to be localized to the cytosol (Hernández et al. 2012), but this study utilized an N-terminus truncated form of the enzyme lacking the cTP. By contrast with *DGAT1*, targeting *DGAT3* via SIGS reduces spore production by 40% (**Fig. 2B**). To confirm the impact of *DGAT3* reduction on powdery mildew asexual reproduction, we obtained a homozygous null mutant in *AtDGAT3*, *dgat3-2* (**Supplemental Fig S2**). *dgat3-2* plants support 21% less spore production than WT (**Fig. 2B**). The larger impact on spore production shown with SIGS rather than null mutants in *DGAT1* and *DGAT3* may be due to genetic compensation through development in the null mutant plants. Neither *dgat1-1* nor *dgat3-2* plants exhibit any obvious developmental or morphological phenotypes.

To determine the localization of AtDGAT3, we cloned the genomic DNA encoding the full length AtDGAT3 sequence and fused 35S to its N-terminus and GFP to its C-terminus. Transient expression of *AtDGAT3-GFP* in *Nicotiana benthamiana* leaves via *Agrobacterium* infiltration results in intense GFP fluorescence that is colocalized with chlorophyll autofluorescence, indicating *DGAT3* is localized to chloroplasts (**Fig. 2C**).

Figure 2D places the tested players in the context of integrated chloroplast-ER lipid metabolism focused on TAG synthesis (Browse et al. 1986; Hölzl and Dörmann 2019; C. Xu, Fan, and Shanklin 2020; Vanhercke et al. 2019), with the addition of AtDGAT3 chloroplast localization. In summary, leaf TAG synthesis to support powdery mildew asexual reproduction occurs using a novel route via DGAT3 in the chloroplast. By contrast, canonical TAG synthesis in the ER via DGAT1 limits spore production.

### Powdery mildew-induced host lipid bodies are present both in the cytosol and chloroplasts

Given our finding that plastidic DGAT3 supports powdery mildew spore production, we performed confocal imaging of infected leaf tissue at 5 and 10 dpi stained with the neutral lipid dye BODIPY505/515 and focused on mesophyll cells underlying powdery mildew feeding structures. As Arabidopsis RPW8.2 is specifically targeted to the fungal extrahaustorial membrane (EHM), we infected Col-0 lines expressing RPW8.2-YFP with *G. orontii* to visualize the haustorium (W. Wang et al. 2009). The haustorium resides in the epidermal cell as depicted in Fig. 1A and is located above three mesophyll cells (Fig. 3A). At the infection site at 5 dpi, abundant lipid droplets are highly localized to the three mesophyll cells right underneath the haustorium (white dashed line in Fig. 3B) and not in distal cells. Almost no lipid bodies are observed in parallel uninfected tissue mesophyll cells. As the infection progresses to 10 dpi, the abundance of lipid bodies increases in the neighboring mesophyll cells. The percent area with fluorescence shows a ∼6-fold increase with infection at 5 dpi and ∼15-fold increase with infection at 10 dpi (Fig. 3C). BODIPY505/515-stained lipid bodies are observed both next to chloroplasts and in the cytoplasm. With closer examination using 3D reconstructions of multiple z-stacked confocal images, we observe that some infection induced lipid bodies are embedded in the chloroplast (Fig. 3D, yellow circled). These chloroplast-embedded lipid bodies (5.2 and 5.9 µm diameter) are similar in size to those that are not embedded; chloroplast adjacent lipid body mean diameter is 3.7µm (n = 8) and cytoplasmic lipid body mean diameter is 3.4 µm (n=5). Together, our results indicate that powdery mildew infection shifts host lipid metabolism to form large storage lipid bodies with some of the lipid bodies inside chloroplasts, others adjacent to or very near the chloroplast, and others in the cytosol.

**Figure 3.**
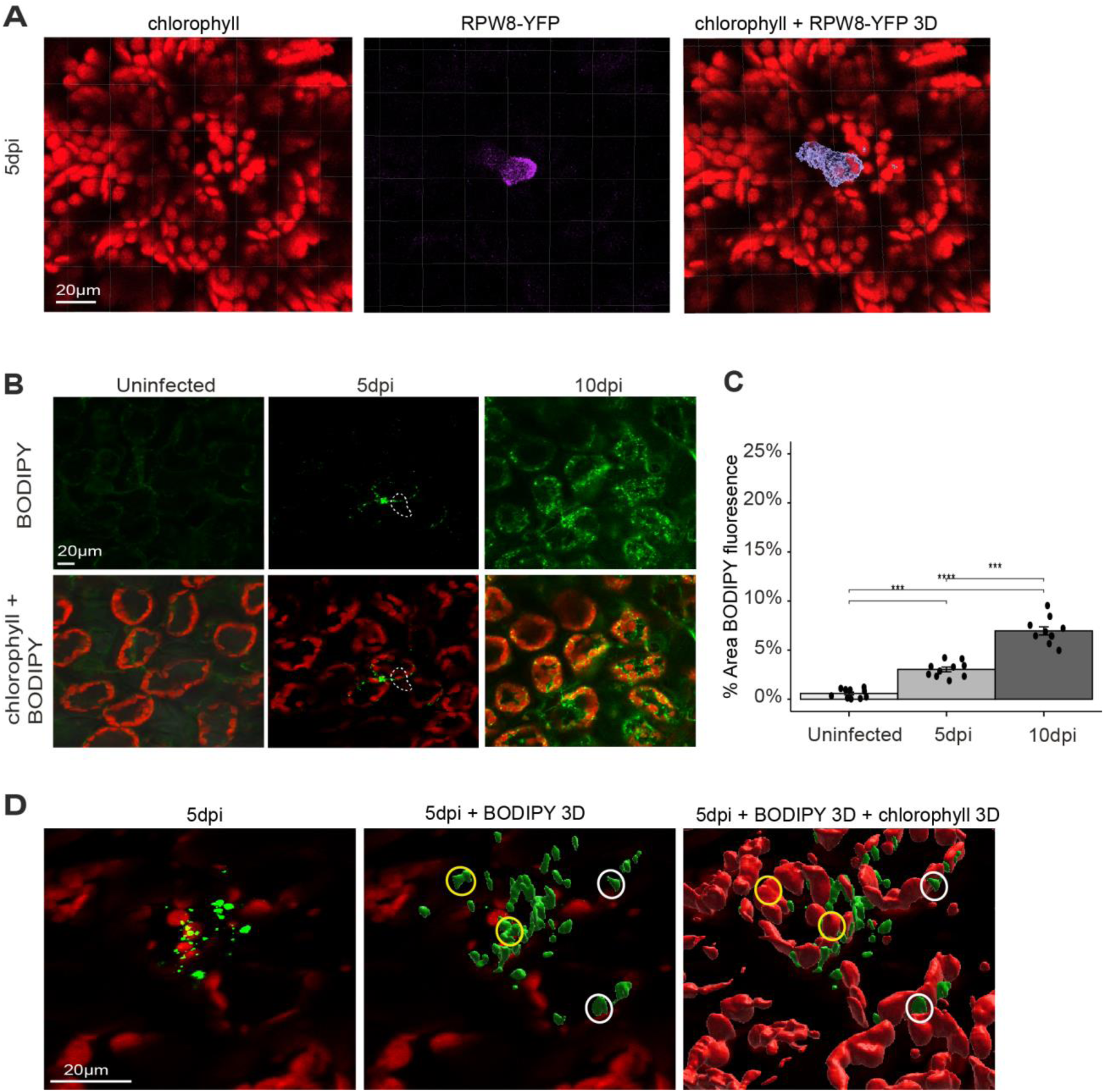
The powdery mildew induces the formation of lipid droplets in the host. A) Representative images of extrahaustorial membrane (EHM) targeted RPW8-YFP showing haustoria in epidermal cell above three mesophyll cells in rosette leaves at 5 days post inoculation (5dpi). B) Representative images of BODIPY 505/515 staining of neutral lipids in mesophyll cell layers of rosette leaves at 5 and 10 dpi. White dash line: position of haustorium in the epidermal cell. C) Percentage of BODIPY fluorescence per image area of 50,000 μm^2^ quantified by Imaris software. Data are mean ± SD of 10 images. Significance is determined by one-way ANOVA. *** p ≤ 0.001, n = 10. D) Representative images of 3D reconstruction of BODIPY fluorescence (green) and chlorophyll fluorescence (red) using Imaris software. Yellow circle: BODIPY fluorescent bodies inside the chloroplast. White circle: BODIPY fluorescence bodies right next to the chloroplast.

### Chloroplast TAG accumulation, but not host defense, is altered in *dgat3-2*

To directly assess whether DGAT3 impacts powdery mildew-induced TAG formation, we performed lipid extractions on 12 dpi leaves and isolated chloroplasts from 12 dpi leaves of *dgat3-2* and WT plants. Using thin layer chromatography (TLC) we find that the TAG content of whole leaf extracts does not differ significantly between *dgat3-2* and WT (Fig. 4A**-B**). However, isolated chloroplast TAG content is reduced by ∼60% in *dgat3-2* compared to WT plants, confirming the role of *DGAT3* in plastidic TAG synthesis. Furthermore, the TAG TLC profile of isolated chloroplasts is enriched in TAGs with a higher Rf than those from whole leaves, overlapping the extra virgin olive oil standard (C18:1 74%, C18:2/3 11%, C16:0 15%).

**Figure 4.**
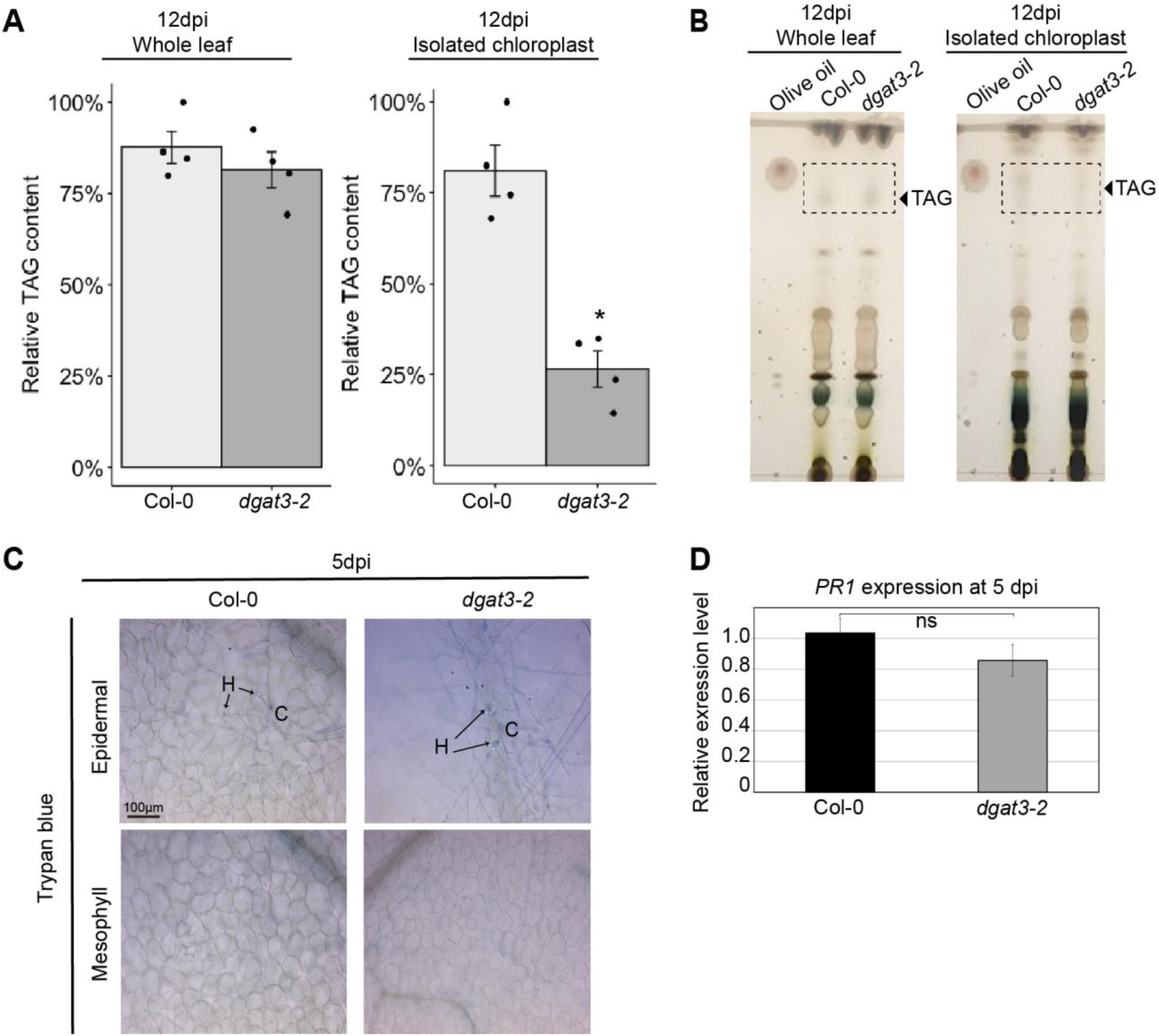
TAG content in infected chloroplasts is decreased in dgat3-2 mutant while defense is not impacted. A) Relative TAG content in whole plants and chloroplasts of Col-0 and *dgat3-2* at 12dpi were quantified by ImageJ software. Data are mean ± SD of 4 biological replicates. Significance is determined by one-way ANOVA, *p ≤ 0.05, n =4. B) Thin-layer chromatography of lipids extracted from either whole plant or isolated chloroplast at 12 dpi. Lipids were visualized with 5% sulfuric acid by charring. C) Trypan blue staining to visualize cell death in Col-0 and *dgat3-2* plants at 5dpi. Top panel, epidermal cell layer. Bottom panel, underlying mesophyll cell layer. H, haustorium; C, germinated conidium. Note that fungal structures are stained slightly by trypan blue. D) Quantitative real-time PCR (qRT-PCR) analysis of *PR1* expression in Col-0 and *dgat3-2* plants at 5dpi normalized to housekeeping gene *ACTIN-2* (± SD, n=3); significance determined using unpaired, two-tailed Student’s T-test. ns= not significantly different at p ≤ 0.05.

Manipulation of plant lipid metabolism can result in altered defense signaling and response including elevated SA responses and/or cell death (Kachroo and Kachroo 2009) that restrict powdery mildew growth and reproduction (e.g. C. A. Frye and Innes 1998; Reuber et al. 1998; Catherine A. Frye, Tang, and Innes 2001). Similar to WT, no cell death is observed in epidermal or mesophyll cells at the powdery mildew infection site of *dgat3-2* plants (Fig. 4C). Moreover, induced *PR-1* expression, a marker of SA-dependent defense responses, does not differ between WT and *dgat3-2* infected leaves (Fig. 4D). Together, these findings suggest the reduction in spore production observed for *dgat3-2* is due to decreased induced plastid TAG production, not increased defense.

### Powdery mildew infection induces the breakdown of thylakoid membrane lipids

Above, we show powdery mildew-induced lipid bodies are associated with chloroplasts (Fig. 3) and plastid-localized AtDGAT3 is a dominant contributor to powdery mildew-induced host TAG synthesis and fungal spore production (**Figs. 2, 4**). As some stresses induce the accumulation of storage lipids at the expense of membrane lipids (Lu et al. 2020; Shiva et al. 2020), we postulated that host chloroplast membranes, dominated by thylakoid membranes, are being disassembled for TAG synthesis in response to infection. We therefore examined the abundance of thylakoid membrane lipids by electrospray ionization (ESI)-MS/MS. Uninfected mature Arabidopsis leaf thylakoid membrane lipids are dominated by monogalactosyldiacylglycerol (MGDG, 42%), digalactosyldiacylglycerol (DGDG, 13%), and phosphatidylglycerol (PG, 10%) (Browse et al. 1989). Powdery mildew-infected (washed) leaves extracted at 12 dpi show that MGDG, DGDG, and PG each decrease by at least 2-fold compared to uninfected leaf controls indicating the breakdown of thylakoid membranes (Fig. 5A**, Supplemental Dataset 2**). Decreased PG, by 60%, was also observed by LC-MS/MS (Fig. 1H).

**Figure 5.**
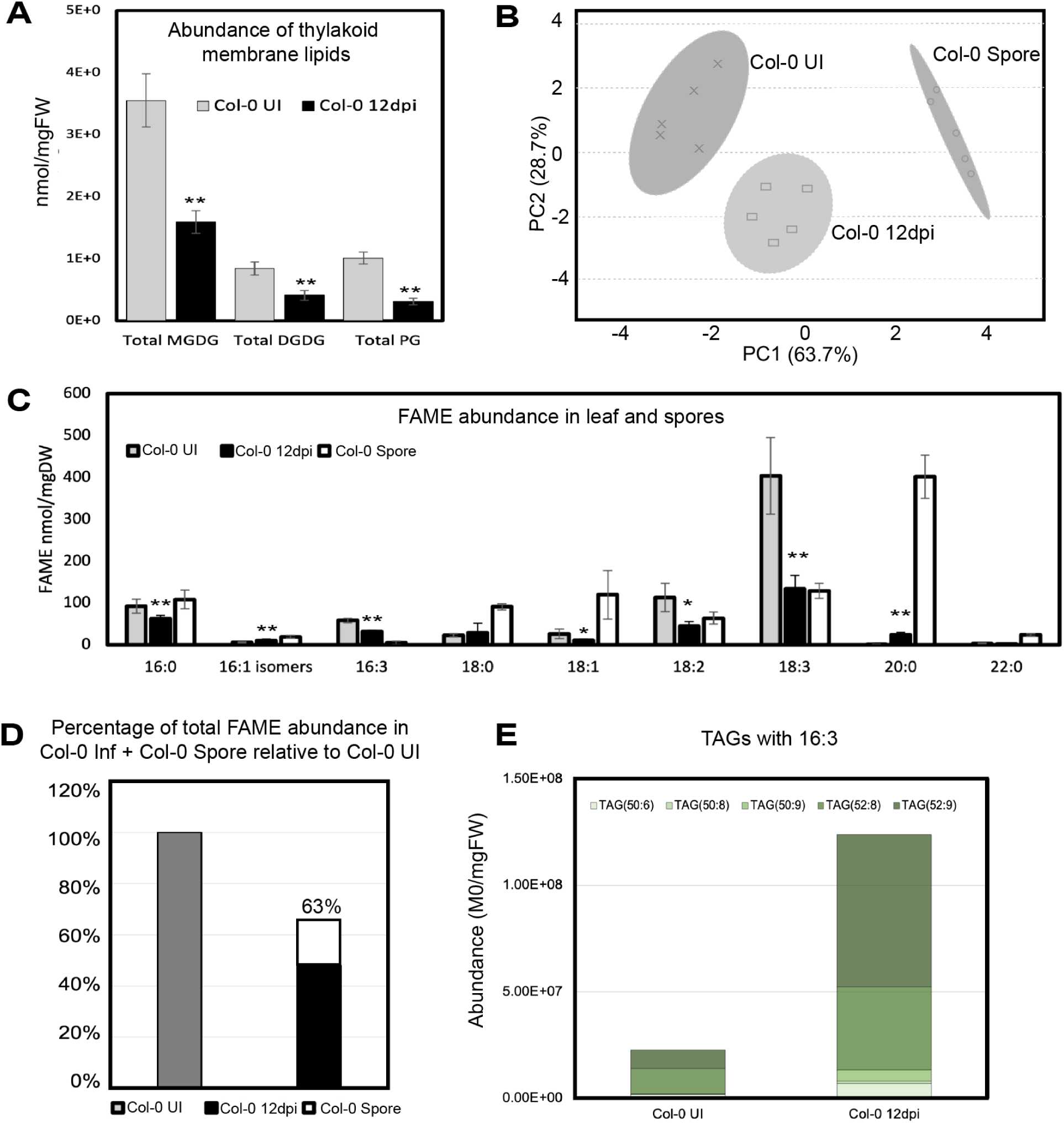
Thylakoid membrane lipids and thylakoid-enriched FAs decrease with infection. A) Abundance of thylakoid membrane lipids (MGDG, monogalactosyldiacylglycerols; DGDG, digalactosyldiacylglycerols; PG, phosphatidylglycerols) in uninfected (UI) and 12dpi leaf lipid extracts (±STD, n= 5). B) Principal component analysis plot based on abundance of FAME species detected (C16-C22) in UI, 12dpi, and spore tissue lipid extracts, n= 5. C) Abundance of FA species detected in the same tissues as in B, normalized to mgDW of that tissue. D) Percentage of total FAME abundance in Col-0 Inf + Col-0 Spore relative to Col-0 UI after conversion of spore data to nmol/mgDW leaf. E) Abundance of TAGs that contain 16:3 in UI and 12 dpi leaf lipid extracts (n=3). Significance between UI and infected leaf samples: 2-tailed T-test *p ≤ 0.05, **p ≤ 0.01.

To understand the change in total FA profiles, lipid extractions were performed on uninfected leaves, washed infected leaves and spore tissues at 12 dpi. Acyl chains were then converted to fatty acid methyl esters (FAMEs) for separation by gas chromatography with flame ionization detection (GC-FID). Principal component analysis (PCA) shows a distinct clustering of the three tissue types according to the ten FA species detected (Fig. 5B). Acyl chains associated with thylakoid membrane lipids, C18:3 (dominant), C18:2, and C16:3 (unique to chloroplast), each decrease by ∼50% in 12 dpi leaves compared with uninfected (Fig. 5C). By contrast, the VLCFA C20:0 increases by ∼20 fold in washed infected leaves. In spore extracts, the VLCFA C20:0 is the dominant species, followed by C18:3, while C18:3 dominates the leaf profiles even after the reduction shown with powdery mildew infection at 12 dpi. By normalizing the spore data to nmol/mgDW leaf (**Supplemental Dataset 2**), we can compare uninfected leaf total FA abundance with that of the washed leaves plus spores. We find 63% of total FA in the uninfected leaves is accounted for in the (washed) infected leaf plus spore (Fig. 5D). Moreover, only the spore C20:0 species abundance is clearly not fully attributed to leaf acquisition as the spore contains 2-fold more C20:0 than the (washed) infected leaf and 36-fold more C20:0 than the uninfected leaf on a leaf normalized basis (**Supplemental Dataset 2**). This also raises the possibility that some of the C20:0 in the (washed) infected leaf FAME samples and LC-MS/MS TAG samples (Fig. 1, **Supplemental Dataset 1**) may be fungal in origin. Only the fungal haustoria is present in the washed leaf samples as all surface structures are removed. Though haustoria make up a small percent of washed leaf sample cells on a cell basis, the haustoria are filled with lipid droplets (Fig. 1) that could include C20:0 remodeled by the fungus. Therefore, it is possible that a portion of the C20:0 in washed leaf analyses is fungal-derived.

As our findings above indicate thylakoid membrane breakdown occurs concurrent with TAG accumulation, we sought to specifically determine whether C16:3 FAs, unique to the thylakoid membrane (Browse et al. 1986), are present in TAGs from 12 dpi leaf lipid extracts (**Supplemental Dataset 1**). Five TAG species were identified as uniquely containing a C16:3 acyl chain, and each of these TAGs also contains at least one C18:3 acyl chain. With infection, these C16:3 containing TAGs increase by 5.4-fold (Fig. 5E). Furthermore, the presence of C16:3 FAs in spores (Fig. 5C, **Supplemental Dataset 2**) indicates fungal acquisition of these thylakoid-derived FAs.

We next sought to examine whether there is an associated change in chloroplast substructures with infection. We examined the ultrastructures of the powdery mildew haustorium and haustorium-associated chloroplasts at 5 dpi via transmission electron microscopy (TEM). The mature haustorium consists of a central haustorium body with peripheral small lobes (Koh et al. 2005). We see abundant electron-dense particles resembling lipid bodies in the haustorium body and lobes and in haustorium-associated chloroplasts (Fig. 6A**-C**). Examination of the haustorium-associated chloroplast in the epidermal cell shows an intact chloroplast outer membrane; however, the thylakoids have considerable loss of grana stacking, indicative of degradation (Fig. 6D).

**Figure 6.**
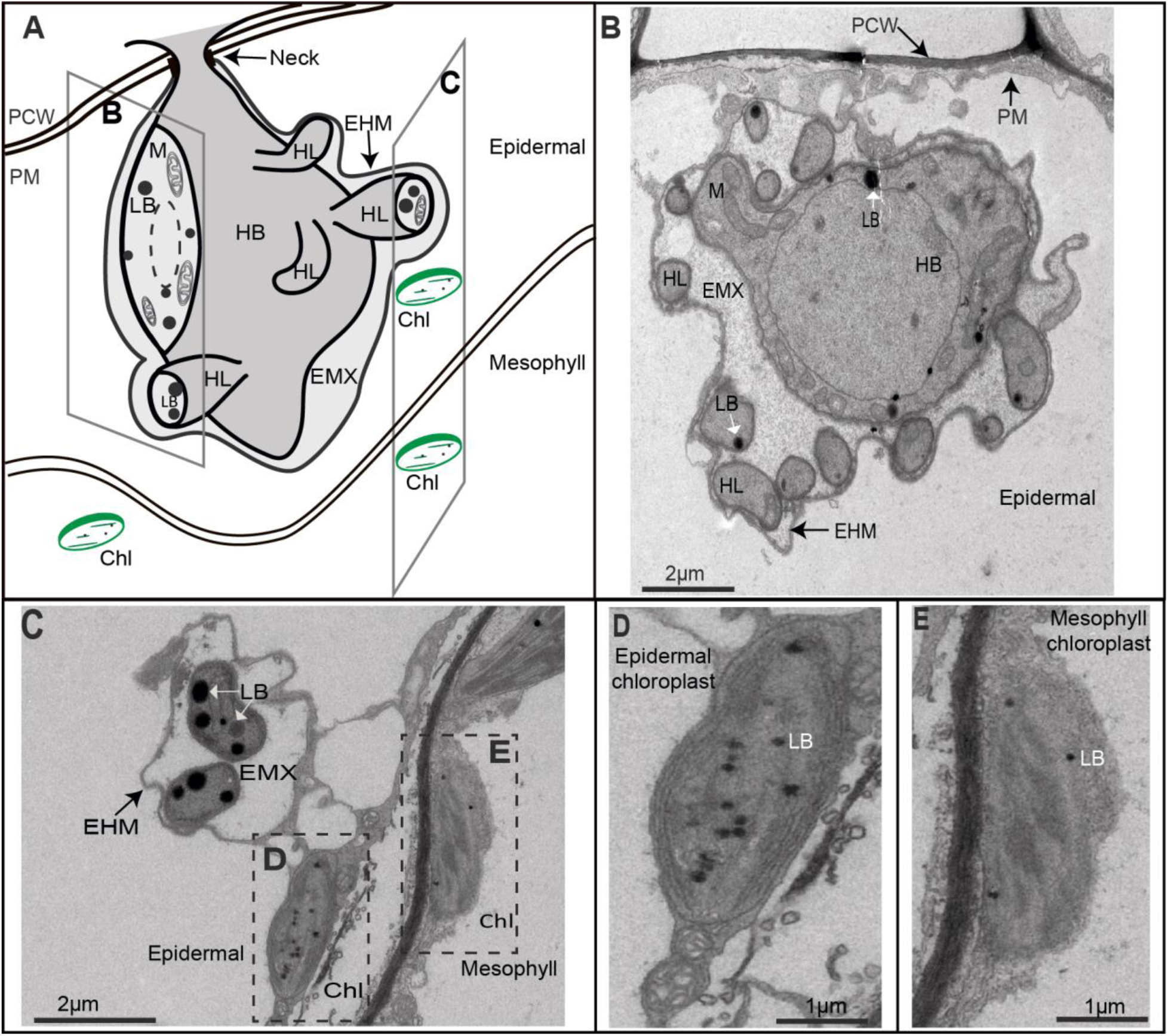
The powdery mildew induces the degradation of host chloroplasts. A) 3D illustration of the powdery mildew haustorium associated with host chloroplasts at 5dpi. B) TEM image of the haustorium. Note this slice does not include the haustorium neck. C) TEM image, slice includes epidermal chloroplast and mesophyll chloroplast associated with the haustorium. D) Zoom-in TEM image, haustorium adjacent epidermal chloroplast. E) Zoom-in TEM image, haustorium adjacent mesophyll chloroplast. Chl, Chloroplast; EHM, Extrahaustorial Membrane; EMX, Extrahaustorial Matix; HB, Haustorium Body; HL, Haustorium Lobe; LB, Lipid Body; M, Mitochondria; PCW, Plant Cell Wall; PM, Plasma Membrane.

Furthermore, the mesophyll chloroplast right underneath the haustorium shows severe degradation, with chloroplast envelope membrane and thylakoid membranes almost totally degraded (Fig. 6E) compared to mesophyll chloroplast from a parallel uninfected leaf (**Supplemental Figure S3**). In addition, no starch is present in this chloroplast. Because the thylakoid membranes are highly degraded in the infected sample, it is difficult to definitively ascertain whether these chloroplast lipid bodies are physically associated with the thylakoid membrane; however, at least one of the three, (Fig. 6E, LB-labeled lipid body), does not appear to be directly attached. Together, our data shows that concurrent with *G.orontii* asexual reproduction (5 dpi+), powdery mildew infection induces the breakdown of host thylakoids, as observed by TEM, with decreased whole leaf thylakoid galactolipids and thylakoid membrane lipid associated FAs.

## DISCUSSION

### Powdery mildew-induced plastidic TAG synthesis utilizes the soluble metalloprotein DGAT3 to promote powdery mildew asexual reproduction

Figure 7 builds on the literature (Browse et al. 1986; Hölzl and Dörmann 2019; C. Xu, Fan, and Shanklin 2020; Vanhercke et al. 2019; C. Xu and Shanklin 2016; Bates 2022) to integrate our findings into a simplified model that shows rewiring of host lipid metabolism by the powdery mildew for TAG synthesis at the expense of thylakoid membranes. In this study, we analyzed the changes in Arabidopsis leaf lipids in response to powdery mildew infection at ≥5dpi concurrent with the formation of spores replete with lipid bodies (Fig. 1B). Despite the highly localized induction of host lipid bodies in mesophyll cells underlying fungal feeding structures (Fig. 3), powdery mildew infected leaves show a 3.5-fold increase in TAG abundance at 12 dpi (Fig. 1D). Localized thylakoid unstacking and degradation (Fig. 6), decreased thylakoid lipids MGDG, DGDG, and PG (Fig. 5A) and decreased thylakoid membrane lipid FAs (Fig. 5C) all suggest TAGs are formed at the expense of thylakoid lipids. This is confirmed by the increase in TAGs containing thylakoid membrane derived acyl chains (18:3 dominant, 18:2, 16:3 unique) (e.g. Fig. 1G**, 5E, Supplemental Dataset 1**) with infection.

**Figure 7.**
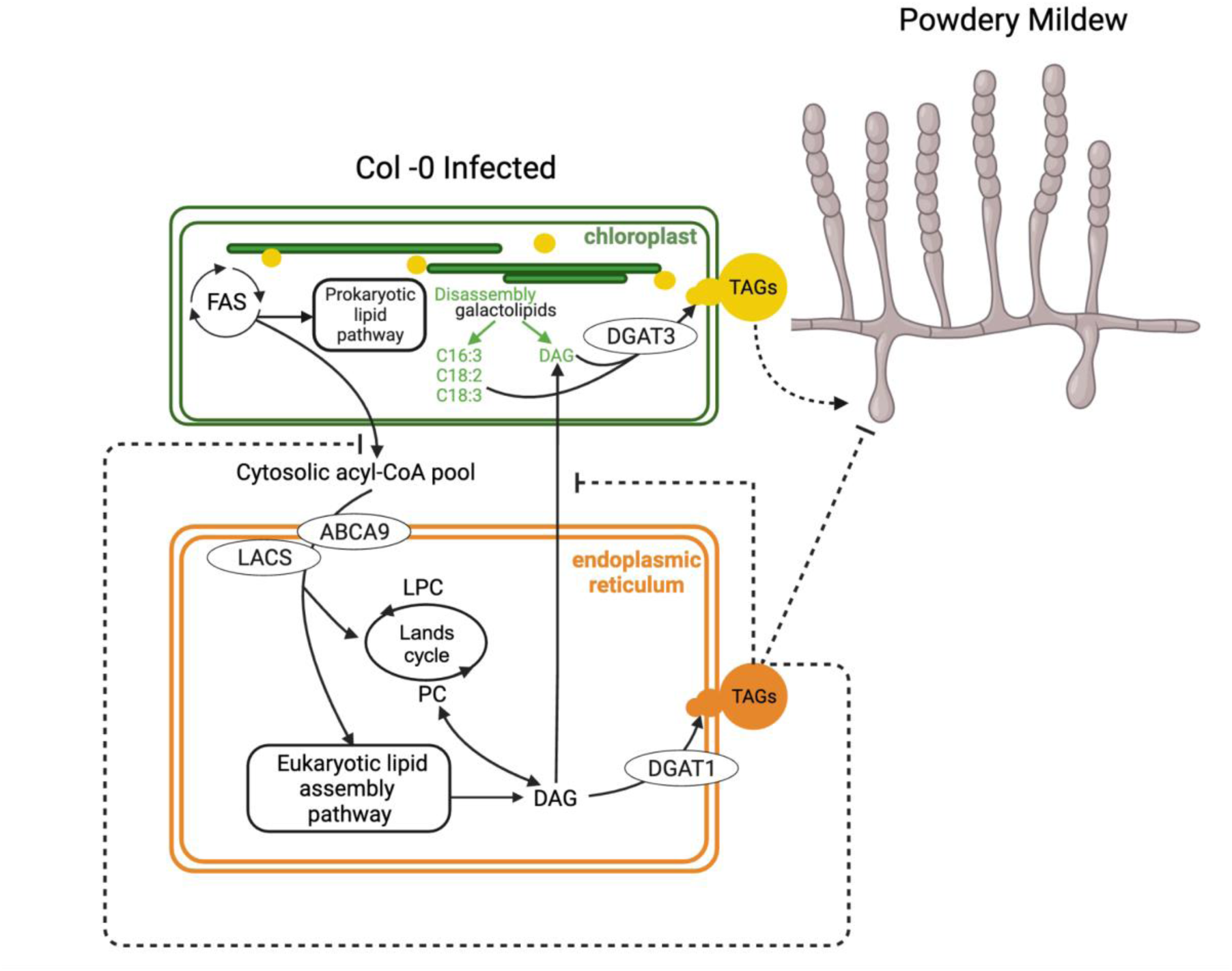
Simplified model for host lipid metabolism rewiring by powdery mildew during its asexual reproduction. Infected Arabidopsis leaves have increased abundance of TAGs and chloroplast-associated and cytosolic lipid bodies concurrent with degradation of thylakoid membranes. In addition, confocal imaging suggests the plastid lipid bodies bleb into the cytosol as shown. Thylakoid lipids and derived fatty acids (FA) decrease with infection and are incorporated into accumulated TAGs. Plastidic TAGs are mostly synthesized by the chloroplast-localized AtDGAT3, which prefers C18:3 and C18:2 substrates, and have a unique profile compared to ER TAGs. Plastid DAGs may be derived from thylakoid membrane breakdown and/or import of DAG/DAG precursors from the ER. Knockdown of *DGAT3* and mutation of *DGAT3* reduced powdery mildew spore production, indicating its function benefits the fungus, likely by supplying energy dense lipids for asexual reproduction and/or providing precursors for a fungal reproductive signal. In contrast, TAGs synthesized via DGAT1 in the ER hinder powdery mildew spore reproduction as assessed using knockouts in the ER fatty acid importer *AtABCA9* and *AtDGAT1*. It is likely that multiple ER LACS activate imported FAs as a knockout in *AtLACS1* alone was insufficient to alter powdery mildew spore production. AtPDAT1 and AtDGAT2, known to use a distinct ER DAG pool for TAG synthesis, do not contribute to powdery mildew asexual reproduction and are not included in the model. ER TAG synthesis via DGAT1 may reduce powdery mildew spore production by limiting substrates for plastidic TAG synthesis and/or supplying lipid droplets that contain TAGs and defensive compounds. Abbreviations: ABCA, ATP-binding cassette A; DAG, diacylglycerol; DGAT, diacylglycerol acyltransferase; FAS, fatty acid synthase complex; LACS, long chain acyl-CoA synthetase; LPC, lysoPC; PC, phosphatidylcholine; PDAT, phospholipid:diacylglycerol acyltransferase; TAGs,triacylglycerols. Dashed lines = proposed.

We further find that the unusual DGAT enzyme, the soluble AtDGAT3 metalloprotein, is localized to the chloroplast (Fig. 2C) and responsible for the bulk (60%) of powdery mildew-induced TAG synthesis in the chloroplast (Fig. 4A**-B**). TLC shows TAGs from chloroplasts isolated from powdery mildew-infected leaves (Fig. 4A**-B**) are enriched in TAGs that run similarly to the extra virgin olive oil standard (85% C18 and 15% C16 FAs). This suggests that the chloroplast TAGs made via AtDGAT3 are enriched for thylakoid-derived acyl chains, as we observe in washed infected whole leaves (Figs. 1G, 5E**, Supplemental Dataset 1**). Furthermore, AtDGAT3 preferentially incorporates C18:3, the dominant FA in thylakoid membranes, and to a lesser extent C18:2 substrates into TAGs <u>(Hernández et al. 2012; Aymé et al. 2018)</u>. It is unclear whether AtDGAT3 may utilize C16:3 as the experimental systems employed by (Hernández et al. 2012; Aymé et al. 2018) had little available C16:3. Powdery mildew spore production is significantly reduced when *AtDGAT3* expression is silenced or when a null mutant in *AtDGAT3* is assessed (Fig. 2B). This reduction in spore production is not associated with a pleiotropic phenotype, enhanced SA defense, and/or cell death in *dgat3-2*. (Fig. 4C**-D**). Therefore, it appears that TAGs synthesized by DGAT3 in the chloroplast at the expense of thylakoid lipids promote powdery mildew asexual reproduction.

At 5 dpi, lipid bodies are observed directly under and in the haustorial complex and are mainly associated with chloroplasts (Figs. 1C, 3, 6). By 10 dpi, chloroplast-associated lipid body accumulation extends to neighboring mesophyll cells underneath the haustorial complex (Fig. 3). As indicated in our model (Fig. 7), 3D reconstructed confocal images suggest chloroplast lipid bodies may then be released into the cytosol for fungal acquisition, as the chloroplast lipid bodies embedded in the chloroplast, adjacent to the chloroplast, and in the cytosol are of similar size (Fig. 3D). It is possible that the induced chloroplast lipid bodies derive (in part) from plastoglobules as our TEM image indicates some lipid bodies in the chloroplast of the mesophyll cell adjacent to the haustorium to be directly associated with the thylakoid membrane (Fig. 6). However, at 5-6 um (Fig. 3D), the lipid bodies are at the top of the size range reported for stress-induced leaf plastoglobules (Arzac, Fernández-Marín, and García-Plazaola 2022; Bouchnak et al. 2023), but common for cytosolic lipid droplets (C. Xu, Fan, and Shanklin 2020).

DGAT3 has not been identified in Arabidopsis plastoglobule proteomics datasets (Ytterberg, Peltier, and van Wijk 2006; Vidi et al. 2006; Lundquist et al. 2012; Espinoza-Corral, Schwenkert, and Lundquist 2021); however, stromal proteins have been identified in plastoglobule subpopulations that also contain thylakoid photosynthetic proteins and lipids but whose membrane varies in composition from that of thylakoid membranes (Ghosh et al. 1994; Smith, Licatalosi, and Thompson 2000). Moreover, plastoglobule blebbing into the stroma and/or release into the cytosol (Ghosh et al. 1994; Springer et al. 2016) has been implicated (C. Xu, Fan, and Shanklin 2020). If the powdery mildew-induced chloroplast lipid bodies derive (in part) from plastoglobules, they may contain the thylakoid membrane-bound phytol ester synthase 1 (PES1) and/or PES2 (Ytterberg, Peltier, and van Wijk 2006; Vidi et al. 2006) which, in addition to phytol ester synthase activity, can synthesize TAGs via DAGs and acyl groups from acyl-CoA (preferred) (Lippold et al. 2012). As 40% of induced chloroplast TAGs remain in *dgat3-2* (Fig. 4A), it is tempting to speculate that in addition to DGAT3, PES1 and/or PES2 also contribute to powdery mildew-induced plastidic TAG synthesis.

How these TAGs directly benefit the powdery mildew remains to be determined. While plastidic TAG catabolism could serve as an immediate energy source, these storage lipids/lipid bodies could also be transported with or without fungal remodeling to the newly developing spores which themselves are filled with lipid bodies containing TAGs (Fig. 1B). The presence of C16:3 acyl chains in spore lipids (Fig. 5C, **Supplemental Dataset 2**) indicates fungal acquisition of the chloroplast TAGs. These spore storage lipids then serve as an energy source to support spore germination and early colonization events prior to haustorium formation (Both et al. 2005). It is also possible that a host-derived lipid may be required for a fungal asexual reproductive signal. For example, in *Aspergillus nidulans* specific endogenous 18:2-derived oxylipins control sporulation versus sexual reproduction (Tsitsigiannis et al. 2004). In the arbuscular mycorrhizal fungi (AMF) - plant host symbiosis, plant derived C16:0 2-MGs are remodeled by the AMF fungus and act both as energy sources (immediate and stored as lipid bodies in spores) and as signals for fungal development, including sporulation (Kameoka et al. 2019). Plastoglobules often contain plant enzymes involved in oxylipin synthesis that could participate in the production of a fungal reproductive signal (Michel, Ponnala, and van Wijk 2021). This could be particularly important for obligate biotrophs such as powdery mildews characterized by missing or incomplete pathways for specialized metabolites as compared to other *Ascomycetes* including *A. nidulans* (Spanu 2012).

### ER-associated TAGs hinder powdery mildew asexual reproduction

To our initial surprise, we found mutants that limit TAG accumulation in the ER exhibit increased powdery mildew spore production (Fig. 2). In Arabidopsis, DGAT1 is responsible for generating TAG from a rapidly produced pool of DAG derived from PC (Regmi et al. 2020). On the other hand, PDAT1 and DGAT2 are reported to use a different and larger pool of DAG, which has a relatively slower turnover (Regmi et al. 2020). Reduced *DGAT1* expression results in enhanced spore production (75% increase, Fig. 2B), while no difference is observed when *PDAT1* or *DGAT2* expression is perturbed (Fig. 2A**,B**). This suggests a rapidly produced pool of DAG from PC available to DGAT1 is used for powdery mildew-induced TAG production in the ER (Fig. 7). We further explored the impact of ABAC9 demonstrated to import FA/acyl-CoA into the ER and found the *abca9-1* mutant supports 50% increased powdery mildew spore production (Fig. 2A). By contrast, the long chain acyl-activating *lacs1-1* mutant, the only ER-localized LACS with enhanced expression at the powdery mildew infection site at 5 dpi (Chandran et al. 2010), did not alter powdery mildew spore production (Fig. 2A). Similarly, a mutant in ER-localized *LACS2* had no impact on powdery mildew growth and reproduction (Tang, Simonich, and Innes 2007). As LACS1, LACS2, LACS4, and LACS8 are all ER-localized (Weng et al. 2010; Zhao et al. 2010; Jessen et al. 2015), it is likely that multiple ER LACS activate imported FAs.

Collectively, our findings indicate induced TAG biosynthesis in the ER via DGAT1 impedes the asexual reproduction of powdery mildew (Fig. 7). AtDGAT1 acyl specificity differs from that of AtDGAT3. C16:0 is the preferred substrate of AtDGAT1, with little activity with C18:2 or C18:3 (Zhou et al. 2013; Aymé et al. 2014). C16:0 is a minor component of thylakoid membrane galactolipids (Browse et al. 1989; Mats X. Andersson, J. Magnus Kjellberg, and Sandelius 2001), consistent with AtDGAT1 use of precursor pools in the ER distinct from those used in the chloroplast by AtDGAT3.

How do ER-synthesized TAGs limit the growth of the biotrophic pathogen? TAGs synthesized at the ER membrane are typically packaged into organelles known as lipid droplets (LDs) that bud from the ER and accumulate in the cytosol (Guzha et al. 2023). Sequestration of these TAGs could be a means of nutrient restriction by the host if these LDs are not accessible to the powdery mildew. Furthermore, given DGAT3-dependent TAG synthesis in the chloroplast supports powdery mildew spore production, it is likely the competing pathway for TAG synthesis in the ER via DGAT1 may divert precursors from the chloroplast pathway (Fig. 7). For example, substrates for plastidic TAG synthesis may be limited by DGAT1 activity pulling plastidic FAs to the ER and/or reducing export of DAG/DAG precursors from the ER to the chloroplast. This competition has been observed in engineered tobacco leaves that accumulate oil at 15% of dry weight (Zhou et al. 2019) and reflects that TAG synthesis drives precursor flux (Bates and Browse 2012).

In addition, LDs not only contain TAGs and sterol esters, but may be sites of specialized biochemistry during stress (Shimada, Hayashi, and Hara-Nishimura 2017). Increased LDs have been observed in leaves infected by the hemi-biotrophic fungus *Colletotrichum higginsanium* and proposed to be sites of phytoalexin synthesis, preventing pathogen spread (Shimada et al. 2014). Furthermore, LDs induced in response to avirulent *Pseudomonas syringae* infection of Arabidopsis leaves were found to contain camalexin biosynthetic enzymes (Fernández-Santos et al. 2020). Genes involved in indole-3-acetaldoxime derived phytoalexin production associated with defense against powdery mildews (Clay et al. 2009; Liu et al. 2016; Hunziker et al. 2020) exhibit enhanced expression at the powdery mildew infection site at 5 dpi (Chandran et al. 2010). This raises the possibility that increased synthesis and/or exposure to defensive specialized metabolites may contribute to the reduction in powdery mildew spore production associated with ER-derived LDs (Fig. 7).

### Powdery mildew infection offers valuable insights into the intricacy of plant lipid metabolism

Although TAGs typically do not accumulate to significant levels in vegetative tissues, TAG accumulation in leaf tissue occurs in response to diverse environmental stresses and leaf senescence (Lu et al. 2020). While a role for AtDGAT3 has not been assessed in response to environmental stresses or leaf senescence, our study shows the important role AtDGAT3 plays in the powdery mildew-host interaction. This indicates AtDGAT3 function should be examined under other conditions, particularly those in which thylakoid disassembly is observed and induced TAGs are enriched in thylakoid-derived FAs, such as response to N limitation and senescence (Kaup, Froese, and Thompson 2002; Gaude et al. 2007; Besagni and Kessler 2013). Our work also argues for tracking cytosolic lipid droplet (LD) origins, as they were previously assumed to be ER-derived. And, the powdery mildew system provides a phenotype (impact on spore production) for distinguishing chloroplast-derived TAGs (via AtDGAT3) from those produced in the ER via AtDGAT1. Whether this translates to other (obligate) plant biotrophs of vegetative tissue remains to be investigated.

As shown by the root colonizing-obligate symbiont AMF, TAGs are only one possible source of lipids for microbial acquisition. AMF manipulate plant root cells to produce 2-MGs for fungal acquisition (Kameoka and Gutjahr 2022). In both systems, localized endoreduplication occurs and is associated with enhanced metabolic capacity that may allow for increased flux to FAs (Wildermuth 2010; Wildermuth et al. 2017). While AMF shifts lipid metabolism to 2-MG production through the use of enzymes specific to AMF host plants, the powdery mildew employs DGAT3, present in almost all land plants (Yan et al. 2018), for chloroplast TAG formation to support asexual reproduction (**Figs. 2, 4**). Therefore, specific host transporters may not be required as they are for AMF 2-MGs. Instead, lipid bodies that originate in the chloroplast have the potential to be directly acquired by the powdery mildew. Similarly, a number of human intracellular pathogens acquire host lipid bodies for their nutrition and development (Vallochi et al. 2018).

### DGAT3, a unique class of DGAT enzyme

The three classes of Arabidopsis DGAT enzymes contain distinct conserved domains and have evolved independently in plants (Yin et al. 2022). The least studied class, DGAT3 enzymes, are unique in that they are soluble metalloproteins, with no transmembrane domain, and a thioredoxin-like ferredoxin domain containing a [2Fe-2S] cluster (Aymé et al. 2018). While AtDGAT3 (Fig. 2C) and *Paeonia rockii* PrDGAT3 (Han et al. 2022) are clearly localized to the chloroplast; other DGAT3 enzymes have been characterized as cytosolic (peanut, (Saha et al. 2006); soybean, (Xue et al. 2022); *Camelina sativa*, (Gao et al. 2021)).

*AtDGAT3* is widely expressed at levels often 10-fold higher than *AtDGAT1* and *AtDGAT2*, with highest expression in the hypocotyl and mature and senescent leaf petioles and stems (Klepikova et al. 2016). Consistent with findings for powdery mildew infection of mature Arabidopsis leaves (Chandran et al. 2009, 2010), *AtDGAT3* is not strongly induced in response to pathogen or abiotic stress, assessed using the Arabidopsis eFP Browser (Winter et al. 2007). As changes in *AtDGAT3* expression are minimal, AtDGAT3 activity may depend on the availability of preferred precursors (e.g. released from thylakoid degradation). Furthermore, AtDGAT3 activity may be regulated by insertion of preformed [2Fe-2S] into the apoprotein in the plastid (Przybyla-Toscano et al. 2018) and by redox.

The availability of [2Fe-2S] clusters, along with maturation factors, could therefore impact DGAT3 metalloprotein levels. We found AtDGAT3 to participate in chloroplast TAG accumulation (Fig. 4) concurrent with thylakoid membrane degradation (**Figs. 5, 6**). When thylakoid membranes are broken down, as we observe in response to powdery mildew, [2Fe-2S] clusters released from thylakoid metalloproteins may be available for insertion into the AtDGAT3 apoprotein. The AtDGAT3 metalloprotein could then help minimize lipotoxicity by converting toxic free fatty acids and DAGs into storage lipids.

Redox state is also likely to regulate the activity of AtDGAT3. Chloroplast redox status, responsive to environmental cues, controls much of chloroplast function including lipid metabolism (Hernández and Cejudo 2021). Ayme et al. (2018) found the AtDGAT3 [2Fe-2S]2+ cluster is stable, while the reduced [2Fe-2S]+ form of the enzyme is rapidly destroyed. When thylakoid membranes are disassembled and/or degraded, reductant generated from oxidative phosphorylation would be decreased and could be insufficient to reduce the AtDGAT3 metallocluster. Similarly, conditions resulting in plastidic oxidative stress (such as high light) could stabilize DGAT3, reducing lipotoxicity.

Engineered plants with increased TAG yield and low input costs for biofuel or specialized chemical applications (Pfleger, Gossing, and Nielsen 2015) could be designed to take advantage of AtDGAT3’s production of TAGs at the expense of thylakoid membranes. The associated TAG profile would be enriched in C18:3 and C18:2 fatty acids desirable for human nutrition (Kumar, Sharma, and Upadhyaya 2016). As shown in Figure 4, the TAGs from isolated chloroplasts infected with powdery mildew appear similar to that of commercial extra virgin olive oil and to be largely attributed to synthesis via AtDGAT3. By contrast the TAGs from infected whole leaves are dominated by TAGs with reduced FA chain length, indicated by the lower Rf, that are likely synthesized in the ER via DGAT1, consistent with its preference for C16:0 (Aymé et al. 2014). Transient expression of *AtDGAT3* or *PrDGAT3* in *N. benthamiana* leaves increases TAG production by ∼2-fold (Hernández et al. 2012; Han et al. 2022), compared to 7-8-fold increase with *AtDGAT1* transient expression (Hernández et al. 2012; Vanhercke et al. 2013). Therefore, in engineered plants, increased flux to plastidic TAG synthesis might be further enhanced by reducing *DGAT1*. In addition, controls over DGAT3 activity and stability would need to be addressed.

Not only does the powdery mildew system allow us to uncover the role of AtDGAT3 in plastid TAG biosynthesis, but it can also be used to dissect key regulators driving flux towards plastid TAG synthesis and lipid body secretion. While the powdery mildew-induced shift in leaf lipid metabolism is highly localized, heavy infection could further increase induced TAG levels from the 3-fold induction observed with the low/moderate levels of infection that facilitate our molecular and microscopic studies. By understanding how mature photosynthetically active leaves switch their metabolism to break down thylakoids to make and secrete storage lipids, higher yields of plant oils could potentially be achieved, than from extracted seeds or fruit. Furthermore, plants suitable for oil production could be expanded and deforestation associated with palm oil plantations could potentially be reduced, facilitating more sustainable and environmentally benign production.

## METHODS

### Plant lines, growth, and powdery mildew infection

*Mutant list:* Seeds of *abca9-1* (SALK_058070, Kim et al. 2013), *lacs1-1* (SALK_127191, Lü et al. 2009), *dgat1-1* (CS3861, Katavic et al. 1995), *dgat3-2* (SALK_112303, **Supplemental Fig. S2**), *pdat1-2* (SALK_065334, Zhang et al. 2009) mutant lines in Col-0 background were obtained from Arabidopsis Biological Resource Center (ABRC) at The Ohio State University. All lines were genotyped to confirm homozygosity, using primers in **Supplemental Table S1**.

Wild type *Arabidopsis thaliana* ecotype Columbia-0 (Col-0) and mutants were grown in SS Metromix200 soil (Sun Gro, Bellevue, WA) in growth chambers at 22°C with 12 h light/dark cycle, 70% relative humidity and PAR of ∼120μmol m^-2^ s^-1^. After stratification at 4°C, alternating Col-0 and mutant seeds were planted in 16.5 cm insert boxes (12 plants/box; 6 boxes/flat). For whole plant spore count phenotyping, boxes of plants were inoculated at 4 weeks by settling tower with a moderate dose of 10-14 dpi conidia from *G. orontii* MGH1 at consistent time of day (Reuber et al. 1998).

### Spray-induced gene silencing (SIGS)

SIGS protocol was adapted from McRae *et.al.* (2023) (McRae et al. 2023). pssRNAit (https://plantgrn.noble.org/pssRNAit/) was used to design an efficient and specific dsRNA for *DGAT1* (AT2G19450), *DGAT2* (AT3G49210), and *DGAT3* (AT1G48300). Templates were amplified (primers in **Supplemental Table S1**) from Col-0 cDNA and prepared for *in vitro* transcription with the HiScribe T7 High Yield RNA Synthesis Kit (New England Biolabs, Ipswich, MA). After purification with Monarch RNA Cleanup Kit (New England Biolabs, Ipswich, MA), RNA was reannealed, quantified and aliquoted in nuclease-free water. 12-15 mature fully expanded Arabidopsis leaves from 4-5 4-week old plants were harvested. Petioles were inserted through a Whatman 1.0 paper overlaid into 1⁄2 MS salts (Research Products International, Prospect, IL), 0.1% 2-(N-morpholino)ethanesulfonic acid (Merck Millipore, Burlington, MA), and 0.8% agar (BD Biosciences, San Jose, CA) in 150 mm plates. Paired plates (with mutant and WT leaves) were placed under the settling tower and infected with 10-14 dpi conidia, as above. 40μg RNA (or nuclease-free mock) was sprayed at 1 hpi and 2 dpi.

### Spore tissue collection and counting

Powdery mildew spore production/mg leaf fresh weight protocol was adapted from Weßling and Panstruga (Weßling and Panstruga 2012). Briefly, at 8-10 dpi, leaves 7-9 from WT and mutant plants in a box, or all 12 leaves from mock and dsRNA-treated plates, were harvested. Spores were washed off leaves by vortexing in 15 mL 0.01% Tween-80 for 30 seconds and filtered through 30μm CellTrics filter (Sysmex America, Lincolnshire, IL) before centrifugation at 4000x*g*. The resulting spore pellet was resuspended in 200-1000μL water. For each sample, nine 1 × 1 mm fields of a Neubauer-improved haemocytometer were counted. For lipid analysis, tissue was immediately frozen and stored until extraction. For spore counting, 3 paired counts of WT and mutant spore suspensions from a box were performed on a Neubauer-improved hemocytometer (Hausser Scientific, Horsham, PA). Spore counts were divided by the fresh weight of the plant tissue to determine spores/mgFW, and then normalized to WT counts. To determine significance, an unpaired, two-tailed Student’s T-test was performed on counts from at least 5 boxes (*p* < 0.05).

### Trypan blue staining

To visualize cell death, leaf tissues were incubated for 16 h at 24°C in the staining solution (2.5 mg/ml trypan blue in lactophenol, lactic acid, glycerol, phenol, water (1:1:1:1)), and two volumes of ethanol were added to this solution. The tissues were cleared in chloral hydrate solution (2.5g/ml chloral hydrate in water) for 16 h at 24°C. Leaf tissues were transferred to 70% glycerol and viewed using the AS Laser Microdissection system microscope (Leica Microsystems, Deerfield, IL). Note that trypan blue also slightly stains fungal structures.

### Reverse transcription (RT)-qPCR analysis

Total RNA from leaves was extracted with RNA using Spectrum (Sigma-Aldrich) Plant Total RNA Kit according to the manufacturer’s protocol. Residual genomic DNA was digested with DNase I (DNaseI, Qiagen). PCR was performed using cDNA using High-Capacity cDNA Reverse Transcription Kit (ThermoFisher Scientific). The gene specific primers for DGAT3 are: P1: 5′-ACCAGAACGGTAGGGTTTCG-3′; P2: 5′-CTAACGTTTGGGCCATCACGAC-3′. Amplification was performed using the following conditions: 95°C for 2 min and 30 cycles of 95°C for 30 s, 60°C for 30 s, and 72°C for 90 s.

To analyze the expression levels of *PR1* in *dgat3-2* and Col-0 with powdery mildew infection, three independently grown biological replicates of two fully expanded leaves (leaves 7-9) at 5 dpi were used for comparison. Tissue was immediately frozen in liquid nitrogen and stored at -80°C until extraction. RNA was extracted using Spectrum (Sigma-Aldrich) Plant Total RNA Kit according to the manufacturer’s protocol. Residual genomic DNA was digested with DNase I (DNaseI, Qiagen). Purity and concentration of RNA was confirmed with Nanodrop-1000 spectrophotometer (ThermoFisher Scientific). Complementary DNA (cDNA) was synthesized from 1μg RNA using High-Capacity cDNA Reverse Transcription Kit (ThermoFisher Scientific). Quantitative real-time PCR (qPCR) experiments were performed in a BioRad CFX96 (BioRad) using the iTaq Universal SYBR Green Supermix (Bio-Rad, USA), following kit instructions. For all genes, thermal cycling started with a 95°C denaturation step for 10 min followed by 40 cycles of denaturation at 95°C for 15 s and annealing at 56°C for 30 s. Each run was finished with melt curve analysis to confirm specificity of amplicon. Three technical replicates were performed for each experimental set. Gene expression (fold change) was calculated normalized to ACTIN2 (At3g18780) as reference gene, and calculated using the Do My qPCR Calculations webtool (http://umrh-bioinfo.clermont.inrae.fr/do_my_qPCRcalc/;) (Tournayre et al. 2019). Primer sequences are provided in **Supplemental Table S1**.

### Golden Gate cloning and transient expression of DGAT3 via Agrobacterium infiltration

The full-length genomic DNA encoding *DGAT3 (AT1G48300)* without stop codon and with removal of an internal restriction site for BsaI was utilized. Two BsaI restriction enzyme sites are added to both 5’ and 3’ end of sequence using PCR primers listed in **Supplemental Table S1**. The sequence was cloned into pICSL22010 plasmid (with C-terminal GFP and CaMV 35S promoter) by Golden Gate cloning. The vector was transformed into *Agrobacterium tumefaciens* GV3101. *A. tumefaciens* transformants were grown in 5 mL liquid LB with appropriate antibiotics overnight at 28°C, pelleted, resuspended in induction media (10 mM MES pH 5.6, 10 mM MgCl2, 150 μM acetosyringone) to an OD600 of 0.4-0.60 for transient expression, and incubated in induction media for approximately 3-4 h before infiltration in *N. benthamiana* leaves. GFP fluorescence was observed at 48-72 hpi by Zeiss LSM710 confocal microscope (Carl Zeiss Inc, White Plains, New York) at the RCNR Biological Imaging Facility, UC Berkeley.

### Confocal imaging

Confocal scanning fluorescence microscopy with a Zeiss LSM710 confocal microscope (Carl Zeiss Inc, White Plains, New York) at the RCNR Biological Imaging Facility, UC Berkeley was utilized to examine fungal haustoria and lipid droplets.

Col-0 lines expressing RPW8.2-YFP under the native promoter were inoculated with *G. orontii* to visualize fungal haustoria (W. Wang et al. 2009). The 3D reconstruction of RPW8.2-YFP was performed using Imaris software. To visualize lipid bodies, tissues were stained with 0.004 mg/mL BODIPY 505/515 and vacuum infiltrated for 10 min before imaging. Excitation of chlorophyll and BODIPY were at 633 and 488 nm, respectively. Emission wavelength for chlorophyll and BODIPY-stained lipid bodies was 647-721 nm and 493-589 nm, respectively. Percent-area of BODIPY fluorescence was quantified using Image J software. The 3D reconstruction of lipid droplets and chloroplasts was performed using Imaris.

### Transmission electron microscopy imaging

Arabidopsis Col-0 4 week old plants were heavily inoculated with *G. orontii*. Leaves were sampled at 5 dpi and cut into 2 × 3-mm sections, fixed in buffer containing 2.5% glutaraldehyde, 2% tween 20, 0.05M sodium cacodylate and 4% formaldehyde in microwave for 2 X 40 s. The fixed tissues were vacuumed for 1 h or as long as possible until they sank to the bottom. The tissues were rinsed three times in 0.05M sodium cacodylate buffer for 10 min. After being transferred into 1% Osmium tetroxide buffer, the tissues were fixed by microwaving for 3 X 1 min, with 15 min vacuum between each microwaving. The samples were dehydrated with a gradient of acetone (35%, 50%, 70%, 80%, 95%, 100%, 100%, 100%) for 10 min each. The tissues were sequentially infiltrated with 20%, 40%, 60% Resin by microwaving (3 min) and rotated overhead for 1 h after each microwaving. The samples were rotated in 80% resin for 16 h. The next day, the samples were rotated in 90% resin 16 h. The samples were embedded in a flat embedding mold and cured in a 60°C oven for 2-3 days. Ultrathin sections were put on mesh nickel grids. After contrast staining, samples were examined and images were acquired with a FEI Tecnai T12 Transmission Electron Microscope at the UC Berkeley Electron Microscopy Laboratory.

### TAGs and Phospholipids via LC-MS/MS

Leaf tissue (leaves 7-9) was harvested at 12 dpi, rapidly weighed, and flash frozen until ready for extraction. After tissue disruption in the bead beater, modified Bligh & Dyer extraction with methanol:chloroform:H_2_0 (1:1:0.9) was performed, 300 μL of chloroform phase was recovered, and dried under nitrogen. The dried extracts were resuspended in 200μL of Isopropanol (IPA):Acetonitrile (ACN):Methanol (MeOH) (3:3:4), and run immediately. Internal standard mixes were used to ensure retention time reproducibility. Samples were run on an Agilent 1290 (Agilent Technologies, Santa Clara, CA) UHPLC connected to a QExactive mass spectrometer (Thermo Fisher Scientific, San Jose, CA) at the DOE Lawrence Berkeley Lab with the following chromatographic method, in both positive and negative mode. Source settings on the MS included auxiliary gas flow of 20 (au), sheath gas flow rate of 55 (au), sweep gas flow of 2 (au), spray voltage of 3 kV (positive and negative ionization modes), and ion transfer tube temperature of 400 °C.

Lipids were run on a reversed phase 50mm x 2.1 mm, 1.8 μm Zorbax RRHD (Rapid Resolution High Definition) C18 column (Agilent Technologies) with a 21 min gradient and 0.4 mL/min flow rate, with 2 μL injections. The mobile phases used were A: 60:40 H_2_O:ACN (60:40) with 5mM ammonium acetate, 0.1% formic acid, and B: IPA:ACN (90:10) with 5mM ammonium acetate (0.2% H2O), 0.1% formic acid. The system was held at 20% B for 1.5 min, followed by an increase to 55% B over 2.5 min, and a subsequent increase to 80% B over 6 min. The system was then held at 80% B for 2 min, before being flushed out with 100% B for 5 min, and re-equilibrated at 20% B over 5 min. The QExactive parameters were as follows: MS resolution was set to 70,000, and data was collected in centroid mode from 80-1200 m/z. MS/MS data was collected at a resolution of 17,500 with a collision energy step gradient of 10, 20, and 30. Lipids were identified by comparing detected vs. theoretical lipid m/z and MS/MS fragmentation patterns, with lipid class and fatty acid composition determined based on characteristic product ions or neutral losses (see **Supplementary Dataset 1**). TAGs were detected in positive ionization mode as [M+NH4]+ adducts, with FA tails determined by neutral loss of ions detected in MS/MS fragmentation spectra. Phospholipids PC, lysoPC, PE, PI, and PG were detected in positive ionization mode as [M+H]+ adducts, with PCs and lysoPCs having a characteristic product ion of 184, PEs a neutral loss of 141, PIs a neutral loss of 260 and PGs a neutral loss of 172 (Murphy 2014). Metabolomics raw data is deposited in the MassIVE data repository (https://massive.ucsd.edu/), accession number MSV000093317 (doi:10.25345/C5N873941). Only lipid classes that have peak heights above the upper bound of the 95% confidence interval of the negative controls are included in **Supplemental Dataset 1** for further analysis.

### Thylakoid Membrane Lipid Analysis

*Tissue harvest and lipid extraction*: Leaves 7-9 were harvested from mock infected and infected plants, washed of spores, frozen in liquid nitrogen, and stored at -80°C until extraction. Extraction was performed following lipase inactivation in 75°C isopropanol for 15 min according to (Devaiah et al. 2006) and electrospray ionization tandem mass spectrometry was performed at the Kansas Lipidomics Research Center Analytical Laboratory (Manhattan, KS) as below.

*Electrospray Ionization Tandem Mass Spectrometry Conditions*: The samples were dissolved in 1 ml chloroform. An aliquot of 10 to 20μl of extract in chloroform was used. Precise amounts of internal standards, obtained and quantified as previously described (Welti et al. 2002), were added in the following quantities (with some small variation in amounts in different batches of internal standards): 0.36 nmol di14:0-PG, 0.36 nmol di24:1-PG, 0.36 nmol 14:0-lysoPG, 0.36 nmol 18:0-lysoPG, 2.01 nmol 16:0-18:0-MGDG, 0.39 nmol di18:0-MGDG, 0.49 nmol 16:0-18:0-DGDG, and 0.71 nmol di18:0-DGDG. Samples were combined with solvents, introduced by continuous infusion into the ESI source on a triple quadrupole MS/MS (API 4000, Applied Biosystems, Foster City, CA), and neutral loss scans were acquired as described by (Shiva et al. 2013).

The background of each spectrum was subtracted, the data were smoothed, and peak areas integrated using a custom script and Applied Biosystems Analyst software. Peaks corresponding to the target lipids in these spectra were identified and the intensities corrected for isotopic overlap. Lipids in each class were quantified in comparison to the two internal standards of that class. The first and typically every 11th set of mass spectra were acquired on the internal standard mixture only. A correction for the reduced response of the mass spectrometer to the galactolipid standards in comparison to its response to the unsaturated leaf galactolipids was applied. To correct for chemical or instrumental noise in the samples, the molar amount of each lipid metabolite detected in the “internal standards only” spectra was subtracted from the molar amount of each metabolite calculated in each set of sample spectra. Finally, the data were corrected for the fraction of the sample analyzed and normalized to the sample leaf dry weight (DW) to produce data in the units nmol/mg DW.

### FAME analysis

Leaves 7-9 were harvested from mock infected and infected plants, washed of spores, frozen in liquid nitrogen, and stored at -80°C until extraction. Extraction was performed following lipase inactivation in 75°C isopropanol for 15 min according to (Devaiah et al. 2006) and FAME analysis was performed by the Kansas Lipidomics Research Center Analytical Laboratory (Manhattan, KS) as below.

Total lipid extracts were spiked with 25 nmol pentadecanoic (C15:0) acid as internal standard. Samples were evaporated under a stream of nitrogen. Samples were resuspended in 1 mL 3 M methanolic hydrochloric acid and heated at 78°C for 30 min. Two mL H2O and 2 mL hexane were added followed by three hexane extractions and then dried down under a stream of nitrogen. Samples were then redissolved in 100 μL hexane and analyzed on GC-FID (Agilent 6890N) after separating sample using a DB-23 capillary column (column length, 60 m; internal diameter, 250 μm; film thickness, 0.25 μm). The carrier was helium gas at a flow rate of 1.5 mL/min. The back inlet was operating at a pressure of 36.01 psi and temperature of 250 °C. The GC oven temperature ramp began with an initial temperature of 150 °C held for 1 min and increased at 25 °C/min to 175 °C. Then the temperature was increased at 4°C/min to 230°C and held at 230°C for 8 min. The total run time was 23.75 min. The flame ionization detector was operated at 260 °C. The hydrogen flow to the detector was 30 mL/min, air flow was 400 mL/min and sampling rate of the FID was 20 Hz. The data were processed using Agilent Chemstation software. As for above, only data with CoV < 0.3 were included. Data are presented as nmol/mg DW of the tissue utilized. Spore nmol/mg DW was multiplied by 0.12995 to indicate the corresponding leaf mg/DW from which the spores were obtained.

### TLC of lipids from isolated chloroplasts and whole leaves

Chloroplast isolation: Leaf tissue from about 40 Arabidopsis plants, infected with *G. orontii* MGH1 at 4-weeks, was harvested at 12 dpi and immediately homogenized by blending for 3x5 s in isolation buffer (30 mM HEPES-KOH pH 8, 0.33M sorbitol, 5 mM MgCl2, 0.1 % [w/v] BSA). The resulting homogenate was briefly filtered through one layer of Miracloth (Chicopee Mills Inc., Milltown, N. J.) Chloroplasts were pelleted with 5-min centrifugation at 1500 g and 4°C, and washed twice with washing buffer (30 mM HEPES-KOH pH 8.0, 0.33M sorbitol). Washed chloroplasts were normalized by chlorophyll concentration and resuspended in an osmotic stress buffer (10 mM Tricine pH 7.9, 1 mM EDTA, 0.6 M sucrose) and stored at −80°C for future analysis.

1-3 mg chloroplasts (normalized by chlorophyll concentrations) or 1-2 g grounded whole leaf tissues from 4-5 week old plants (normalized by fresh weight) for infected samples at 12 dpi were sonicated with 4 pulses of 10 sec and 20% wattage (Model VCX 130, Sonics & Materials INC, Newtown, CT). 1 mL of 2:1 Chloroform Methanol (v/v) with 0.01%BHT was added and placed on a vortex for 5 min. 266 µL of 0.73% (w/v) NaCl solution was added, and the mixture was inverted 5-6 times to mix. Samples were then centrifuged for 5 min at 10,000 × g. The lower, solvent phase was used and dried under an N_2_ stream and resuspended in 20 µL chloroform. In total, 10 µL of the concentrated lipid extract was loaded onto a clean silica TLC plate (MilliporeSigma™ TLC Silica Gel 60 F254: 25 Glass plates, M1057150001) and developed hexane:diethyl ether:glacial acetic acid (91:39:1.3) for 30 min. Lipids were visualized by sulfuric acid spray and charring (25% H2SO4 in 50% ethanol, 135 °C for 10 min). Trader Giotto’s extra virgin olive oil (0.01ug loaded) was used as a standard. TLC was conducted for four separate experiments, each serving as a biological replicate. Relative TAG content analysis was performed using ImageJ software.

## ACCESSION NUMBERS

*ABCA9* (AT5G61730)*, LACS1* (AT2G47240)*, PDAT1* (AT5G13640)*, DGAT1* (AT2G19450), *DGAT2* (AT3G51520)*, DGAT3* (AT1G48300), *PR1* (At2g14610), MassIVE data repository (https://massive.ucsd.edu/) accession number MSV000093317.

## SUPPLEMENTAL MATERIALS

**Supplemental Table S1.** Genotyping, cloning, and SIGS dsRNA template primers used for this work.

**Supplemental Figure S1.** Abundance of TAG species detected in infected leaves at 12 dpi, compared to uninfected leaves.

**Supplemental Figure S2.** Identification of *dgat3-2* (SALK_112303) mutant.

**Supplemental Figure S3.** Transmission electron microscopy image of mesophyll chloroplast from uninfected Arabidopsis leaf.

**Supplemental Dataset 1: LC-MS/MS analysis.**

**Supplemental Dataset 2: FAME and ESI-MS/MS analysis.**

## ACKNOWLEDGEMENT

This work was supported by National Science Foundation (NSF) MCB-1617020 and PFI-1919244, and USDA National Institute of Food and Agriculture, Hatch Project Accession Number 1016994 awards to MCW. HX also received a NRAEF Viticulture and Enology Scholarship. JS was supported through NSF MCB-1617020, with TN as co-PI. KL acknowledges support from the US Department of Energy Joint Genome Institute (https://ror.org/04xm1d337; operated under Contract No. DE-AC02-05CH11231 to Lawrence Berkeley National Laboratory). We thank Dr. Denise Schichnes at the RCNR Biological Imaging Facility and the staff of the Electron Microscope Laboratory at the University of California Berkeley for their assistance. Microscopy imaging reported in this publication was supported in part by the National Institutes of Health S10 program under award number 1S10RR026866-01. The thylakoid lipid and FAME analyses described in this work were performed at the Kansas Lipidomics Research Center Analytical Laboratory (Manhattan, KS) by Mary Roth and Libin Yao. Instrument acquisition and lipidomics method development were supported by the National Science Foundation (including support from the Major Research Instrumentation program; most recent award DBI-1726527), K-IDeA Networks of Biomedical Research Excellence (INBRE) of National Institute of Health (P20GM103418), USDA National Institute of Food and Agriculture (Hatch/Multi-State project 1013013), and Kansas State University. We thank Dr. Shunyuan Xiao (University of Maryland, College Park) for RPW8-YFP Arabidopsis seeds, Dr. Ksenia Krasileva (UC Berkeley) for plasmid pICSL22010, and Dr. Krishna Niyogi (UC Berkeley) and Dr. Peter Dörmann (University of Bonn, Germany) for critical review of the manuscript.

## AUTHOR CONTRIBUTIONS

JJ, HX and MCW planned and designed the research. Spore counting of mutant and WT plants was performed by JJ and RM. JJ prepared samples for lipid analyses. HX performed DGAT3 cloning and characterization. Confocal imaging was done by HX, and TEM by HX and JJ. JS, KL, and TN performed the LC-MS/MS TAG and PL analyses. JJ, HX and MCW wrote the manuscript. All authors contributed to the reviewing of the manuscript.

## REFERENCES

1. Abood, J. K., and Dorothy M. Lösel. 1989. “Effects of Powdery Mildew Infection on the Lipid Metabolism of Cucumber.” In Biological Role of Plant Lipids, edited by Péter A. Biacs, Katalin Gruiz, and Tibor Kremmer, 597–601. Boston, MA: Springer US.

2. Arzac, Miren I., Beatriz Fernández-Marín, and José I. García-Plazaola. 2022. “More than Just Lipid Balls: Quantitative Analysis of Plastoglobule Attributes and Their Stress-Related Responses.” Planta 255 (3): 62.

3. Atella, Georgia C., Paula R. Bittencourt-Cunha, Rodrigo D. Nunes, Mohammed Shahabuddin, and Mário A. C. Silva-Neto. 2009. “The Major Insect Lipoprotein Is a Lipid Source to Mosquito Stages of Malaria Parasite.” Acta Tropica 109 (2): 159–62.

4. Aymé, Laure, Simon Arragain, Michel Canonge, Sébastien Baud, Nadia Touati, Ornella Bimai, Franjo Jagic, et al. 2018. “Arabidopsis Thaliana DGAT3 Is a [2Fe-2S] Protein Involved in TAG Biosynthesis.” Scientific Reports 8 (1): 1–10.

5. Aymé, Laure, Sébastien Baud, Bertrand Dubreucq, Florent Joffre, and Thierry Char 2014. “Function and Localization of the Arabidopsis Thaliana Diacylglycerol Acyltransferase DGAT2 Expressed in Yeast.” PloS One 9 (3): e92237.

6. Bates, Philip D. 2022. “Chapter Six - The Plant Lipid Metabolic Network for Assembly of Diverse Triacylglycerol Molecular Species.” In Advances in Botanical Research, edited by Fabrice Rébeillé and Eric Maréchal, 101:225–52. Academic Press.

7. Bates, Philip D., and John Browse. 2012. “The Significance of Different Diacylgycerol Synthesis Pathways on Plant Oil Composition and Bioengineering.” Frontiers in Plant Science 3 (July): 147.

8. Baud, Sébastien, Bertrand Dubreucq, Martine Miquel, Christine Rochat, and Loïc Lepiniec. 2008. “Storage Reserve Accumulation in Arabidopsis: Metabolic and Developmental Control of Seed Filling.” The Arabidopsis Book / American Society of Plant Biologists 6 (July): e0113.

9. Besagni, Céline, and Felix Kessler. 2013. “A Mechanism Implicating Plastoglobules in Thylakoid Disassembly during Senescence and Nitrogen Starvation.” Planta 237 (2): 463– 70.

10. Both, Maike, Michael Csukai, Michael P. H. Stumpf, and Pietro D. Spanu. 2005. “Gene Expression Profiles of Blumeria Graminis Indicate Dynamic Changes to Primary Metabolism during Development of an Obligate Biotrophic Pathogen.” The Plant Cell 17(7): 2107–22.

11. Bouchnak, Imen, Denis Coulon, Vincent Salis, Sabine D’Andréa, and Claire Bréhélin. 2023. “Lipid Droplets Are Versatile Organelles Involved in Plant Development and Plant Response to Environmental Changes.” Frontiers in Plant Science 14 (June): 1193905.

12. Browse, J., L. Kunst, S. Anderson, S. Hugly, and C. Somerville. 1989. “A Mutant of Arabidopsis Deficient in the Chloroplast 16:1/18:1 Desaturase.” Plant Physiology 90 (2): 522–29.

13. Browse, J., N. Warwick, C. R. Somerville, and C. R. Slack. 1986. “Fluxes through the Prokaryotic and Eukaryotic Pathways of Lipid Synthesis in the ‘16:3’ Plant Arabidopsis Thaliana.” Biochemical Journal 235 (1): 25–31.

14. Cavaco, Ana Rita, Ana Rita Matos, and Andreia Figueiredo. 2021. “Speaking the Language of Lipids: The Cross-Talk between Plants and Pathogens in Defence and Disease.” Cellular and Molecular Life Sciences: CMLS 78 (9): 4399–4415.

15. Chandran, Divya, Noriko Inada, Greg Hather, Christiane K. Kleindt, and Mary C. Wildermuth. 2010. “Laser Microdissection of Arabidopsis Cells at the Powdery Mildew Infection Site Reveals Site-Specific Processes and Regulators.” Proceedings of the National Academy of Sciences of the United States of America 107 (1): 460–65.

16. Chandran, Divya, Yu Chuan Tai, Gregory Hather, Julia Dewdney, Carine Denoux, Diane G. Burgess, Frederick M. Ausubel, Terence P. Speed, and Mary C. Wildermuth. 2009. “Temporal Global Expression Data Reveal Known and Novel Salicylate-Impacted Processes and Regulators Mediating Powdery Mildew Growth and Reproduction on Arabidopsis.” Plant Physiology 149 (3): 1435–51.

17. Clark, Joanna I. M., and J. L. Hall. 1998. “Solute Transport into Healthy and Powdery Mildew-Infected Leaves of Pea and Uptake by Powdery Mildew Mycelium.” The New Phytologist 140 (2): 261–69.

18. Clay, Nicole K., Adewale M. Adio, Carine Denoux, Georg Jander, and Frederick M. Ausubel. 2009. “Glucosinolate Metabolites Required for an Arabidopsis Innate Immune Response.” Science 323 (5910): 95–101.

19. Costa, G., M. Gildenhard, M. Eldering, R. L. Lindquist, A. E. Hauser, R. Sauerwein, C. Goosmann, V. Brinkmann, P. Carrillo-Bustamante, and E. A. Levashina. 2018. “Non-Competitive Resource Exploitation within Mosquito Shapes within-Host Malaria Infectivity and Virulence.” Nature Communications 9 (1): 1–11.

20. Dahlqvist, Anders, Ulf Ståhl, Marit Lenman, Antoni Banas, Michael Lee, Line Sandager, Hans Ronne, and Sten Stymne. 2000. “Phospholipid:diacylglycerol Acyltransferase: An Enzyme That Catalyzes the Acyl-CoA-Independent Formation of Triacylglycerol in Yeast and Plants.” Proceedings of the National Academy of Sciences 97 (12): 6487–92.

21. Devaiah, Shivakumar Pattada, Mary R. Roth, Ethan Baughman, Maoyin Li, Pamela Tamura, Richard Jeannotte, Ruth Welti, and Xuemin Wang. 2006. “Quantitative Profiling of Polar Glycerolipid Species from Organs of Wild-Type Arabidopsis and a Phospholipase Dalpha1 Knockout Mutant.” Phytochemistry 67 (17): 1907–24.

22. Espinoza-Corral, Roberto, Serena Schwenkert, and Peter K. Lundquist. 2021. “Molecular Changes of Arabidopsis Thaliana Plastoglobules Facilitate Thylakoid Membrane Remodeling under High Light Stress.” The Plant Journal: For Cell and Molecular Biology 106 (6): 1571–87.

23. Fan, Jilian, Chengshi Yan, and Changcheng Xu. 2013. “Phospholipid:diacylglycerol Acyltransferase-Mediated Triacylglycerol Biosynthesis Is Crucial for Protection against Fatty Acid-Induced Cell Death in Growing Tissues of Arabidopsis.” The Plant Journal: For Cell and Molecular Biology 76 (6): 930–42.

24. Fernández-Santos, Rubén, Yovanny Izquierdo, Ana López, Luis Muñiz, Marta Martínez, Tomás Cascón, Mats Hamberg, and Carmen Castresana. 2020. “Protein Profiles of Lipid Droplets during the Hypersensitive Defense Response of Arabidopsis against Pseudomonas Infection.” Plant & Cell Physiology 61 (6): 1144–57.

25. Fotopoulos, Vasileios, Martin J. Gilbert, Jon K. Pittman, Alison C. Marvier, Aram J. Buchanan, Norbert Sauer, J. L. Hall, and Lorraine E. Williams. 2003. “The Monosaccharide Transporter Gene, AtSTP4, and the Cell-Wall Invertase, Atbetafruct1, Are Induced in Arabidopsis during Infection with the Fungal Biotroph Erysiphe Cichoracearum.” Plant Physiology 132 (2): 821–29.

26. Frye, C. A., and R. W. Innes. 1998. “An Arabidopsis Mutant with Enhanced Resistance to Powdery Mildew.” The Plant Cell 10 (6): 947–56.

27. Frye, Catherine A., Dingzhong Tang, and Roger W. Innes. 2001. “Negative Regulation of Defense Responses in Plants by a Conserved MAPKK Kinase.” Proceedings of the National Academy of Sciences 98 (1): 373–78.

28. Gao, Huiling, Yu Gao, Fei Zhang, Baoling Liu, Chunli Ji, Jinai Xue, Lixia Yuan, and Runzhi Li. 2021. “Functional Characterization of an Novel Acyl-CoA:diacylglycerol Acyltransferase 3-3 (CsDGAT3-3) Gene from Camelina Sativa.” Plant Science: An International Journal of Experimental Plant Biology 303 (February): 110752.

29. Gaude, Nicole, Claire Bréhélin, Gilbert Tischendorf, Felix Kessler, and Peter Dörmann. 2007. “Nitrogen Deficiency in Arabidopsis Affects Galactolipid Composition and Gene Expression and Results in Accumulation of Fatty Acid Phytyl Esters.” The Plant Journal: For Cell and Molecular Biology 49 (4): 729–39.

30. Ghosh, S., K. A. Hudak, E. B. Dumbroff, and J. E. Thompson. 1994. “Release of Photosynthetic Protein Catabolites by Blebbing from Thylakoids.” Plant Physiology 106 (4): 1547–53.

31. Glawe, Dean A. 2008. “The Powdery Mildews: A Review of the World’s Most Familiar (yet Poorly Known) Plant Pathogens.” Annual Review of Phytopathology 46: 27–51.

32. Guzha, Athanas, Payton Whitehead, Till Ischebeck, and Kent D. Chapman. 2023. “Lipid Droplets: Packing Hydrophobic Molecules Within the Aqueous Cytoplasm.” Annual Review of Plant Biology 74 (1): 195–223.

33. Han, Longyan, Yuhui Zhai, Yumeng Wang, Xiangrui Shi, Yanfeng Xu, Shuguang Gao, Man Zhang, Jianrang Luo, and Qingyu Zhang. 2022. “Diacylglycerol Acyltransferase 3(DGAT3) Is Responsible for the Biosynthesis of Unsaturated Fatty Acids in Vegetative Organs of Paeonia Rockii.” International Journal of Molecular Sciences 23 (22). 10.3390/ijms232214390.

34. Hernández, M. Luisa, and Francisco Javier Cejudo. 2021. “Chloroplast Lipids Metabolism and Function. A Redox Perspective.” Frontiers in Plant Science 12 (August): 712022.

35. Hernández, M. Luisa, Lynne Whitehead, Zhesi He, Valeria Gazda, Alison Gilday, Ekaterina Kozhevnikova, Fabián E. Vaistij, Tony R. Larson, and Ian A. Graham. 2012. “A Cytosolic Acyltransferase Contributes to Triacylglycerol Synthesis in Sucrose-Rescued Arabidopsis Seed Oil Catabolism Mutants.” Plant Physiology 160 (1): 215–25.

36. Hölzl, Georg, and Peter Dörmann. 2019. “Chloroplast Lipids and Their Biosynthesis.” Annual Review of Plant Biology 70 (April): 51–81.

37. Hunziker, Pascal, Hassan Ghareeb, Lena Wagenknecht, Christoph Crocoll, Barbara Ann Halkier, Volker Lipka, and Alexander Schulz. 2020. “De Novo Indol-3-Ylmethyl Glucosinolate Biosynthesis, and Not Long-Distance Transport, Contributes to Defence of Arabidopsis against Powdery Mildew.” Plant, Cell & Environment 43 (6): 1571–83.

38. Jessen, Dirk, Charlotte Roth, Marcel Wiermer, and Martin Fulda. 2015. “Two Activities of Long-Chain Acyl-Coenzyme A Synthetase Are Involved in Lipid Trafficking between the Endoplasmic Reticulum and the Plastid in Arabidopsis.” Plant Physiology 167 (2): 351–66.

39. Jiang, Yina, Wanxiao Wang, Qiujin Xie, Na Liu, Lixia Liu, Dapeng Wang, Xiaowei Zhang, et al. 2017. “Plants Transfer Lipids to Sustain Colonization by Mutualistic Mycorrhizal and Parasitic Fungi.” Science 356 (6343): 1172–75.

40. Kachroo, Aardra, and Pradeep Kachroo. 2009. “Fatty Acid–Derived Signals in Plant Defense.” Annual Review of Phytopathology 47 (1): 153–76.

41. Kameoka, Hiromu, and Caroline Gutjahr. 2022. “Functions of Lipids in Development and Reproduction of Arbuscular Mycorrhizal Fungi.” Plant & Cell Physiology 63 (10): 1356– 65.

42. Kameoka, Hiromu, Taro Maeda, Nao Okuma, and Masayoshi Kawaguchi. 2019. “Structure-Specific Regulation of Nutrient Transport and Metabolism in Arbuscular Mycorrhizal Fungi.” Plant & Cell Physiology 60 (10): 2272–81.

43. Katavic, V., D. W. Reed, D. C. Taylor, E. M. Giblin, D. L. Barton, J. Zou, S. L. Mackenzie, P. S. Covello, and L. Kunst. 1995. “Alteration of Seed Fatty Acid Composition by an Ethyl Methanesulfonate-Induced Mutation in Arabidopsis Thaliana Affecting Diacylglycerol Acyltransferase Activity.” Plant Physiology 108 (1): 399–409.

44. Kaup, Marianne T., Carol D. Froese, and John E. Thompson. 2002. “A Role for Diacylglycerol Acyltransferase during Leaf Senescence.” Plant Physiology 129 (4): 1616–26.

45. Kim, Sangwoo, Yasuyo Yamaoka, Hirofumi Ono, Hanul Kim, Donghwan Shim, Masayoshi Maeshima, Enrico Martinoia, Edgar B. Cahoon, Ikuo Nishida, and Youngsook Lee. 2013. “AtABCA9 Transporter Supplies Fatty Acids for Lipid Synthesis to the Endoplasmic Reticulum.” Proceedings of the National Academy of Sciences of the United States of America 110 (2): 773–78.

46. Klepikova, Anna V., Artem S. Kasianov, Evgeny S. Gerasimov, Maria D. Logacheva, and Aleksey A. Penin. 2016. “A High Resolution Map of the Arabidopsis Thaliana Developmental Transcriptome Based on RNA-Seq Profiling.” The Plant Journal: For Cell and Molecular Biology 88 (6): 1058–70.

47. Koh, Serry, Aurélie André, Herb Edwards, David Ehrhardt, and Shauna Somerville. 2005. “Arabidopsis Thaliana Subcellular Responses to Compatible Erysiphe Cichoracearum Infections.” The Plant Journal: For Cell and Molecular Biology 44 (3): 516–29.

48. Kumar, Aruna, Aarti Sharma, and Kailash C. Upadhyaya. 2016. “Vegetable Oil: Nutritional and Industrial Perspective.” Current Genomics 17 (3): 230–40.

49. Liang, Peng, Songyu Liu, Feng Xu, Shuqin Jiang, Jun Yan, Qiguang He, Wenbo Liu, et al. 2018. “Powdery Mildews Are Characterized by Contracted Carbohydrate Metabolism and Diverse Effectors to Adapt to Obligate Biotrophic Lifestyle.” Frontiers in Microbiology 9 (December): 3160.

50. Lippold, Felix, Katharina vom Dorp, Marion Abraham, Georg Hölzl, Vera Wewer, Jenny Lindberg Yilmaz, Ida Lager, et al. 2012. “Fatty Acid Phytyl Ester Synthesis in Chloroplasts of Arabidopsis.” The Plant Cell 24 (5): 2001–14.

51. Liu, Simu, Lisa M. Bartnikas, Sigrid M. Volko, Frederick M. Ausubel, and Dingzhong Tang. 2016. “Mutation of the Glucosinolate Biosynthesis Enzyme Cytochrome P450 83A1 Monooxygenase Increases Camalexin Accumulation and Powdery Mildew Resistance.” Frontiers in Plant Science 7 (March): 227.

52. Luginbuehl, Leonie H., Guillaume N. Menard, Smita Kurup, Harrie Van Erp, Guru V. Radhakrishnan, Andrew Breakspear, Giles E. D. Oldroyd, and Peter J. Eastmond. 2017. “Fatty Acids in Arbuscular Mycorrhizal Fungi Are Synthesized by the Host Plant.” Science 356 (6343): 1175–78.

53. Lu, Junhao, Yang Xu, Juli Wang, Stacy D. Singer, and Guanqun Chen. 2020. “The Role of Triacylglycerol in Plant Stress Response.” Plants 9 (4). 10.3390/plants9040472.

54. Lundquist, Peter K., Anton Poliakov, Nazmul H. Bhuiyan, Boris Zybailov, Qi Sun, and Klaas J. van Wijk. 2012. “The Functional Network of the Arabidopsis Plastoglobule Proteome Based on Quantitative Proteomics and Genome-Wide Coexpression Analysis.” Plant Physiology 158 (3): 1172–92.

55. Lü, Shiyou, Tao Song, Dylan K. Kosma, Eugene P. Parsons, Owen Rowland, and Matthew A. Jenks. 2009. “Arabidopsis CER8 Encodes LONG-CHAIN ACYL-COA SYNTHETASE 1 (LACS1) That Has Overlapping Functions with LACS2 in Plant Wax and Cutin Synthesis.” The Plant Journal: For Cell and Molecular Biology 59 (4): 553–64.

56. MacLean, Allyson M., Armando Bravo, and Maria J. Harrison. 2017. “Plant Signaling and Metabolic Pathways Enabling Arbuscular Mycorrhizal Symbiosis.” The Plant Cell 29 (10): 2319–35.

57. Mats X. Andersson, J. Magnus Kjellberg, and Anna Stina Sandelius. 2001. “Chloroplast Biogenesis. Regulation of Lipid Transport to the Thylakoid in Chloroplasts Isolated from Expanding and Fully Expanded Leaves of Pea.” Plant Physiology 127 (1): 184–93.

58. McRae, Amanda G., Jyoti Taneja, Kathleen Yee, Xinyi Shi, Sajeet Haridas, Kurt LaButti, Vasanth Singan, Igor V. Grigoriev, and Mary C. Wildermuth. 2023. “Spray-Induced Gene Silencing to Identify Powdery Mildew Gene Targets and Processes for Powdery Mildew Control.” Molecular Plant Pathology 24 (9): 1168–83.

59. Micali, Cristina, Katharina Göllner, Matt Humphry, Chiara Consonni, and Ralph Panstruga. 2008. “The Powdery Mildew Disease of Arabidopsis: A Paradigm for the Interaction between Plants and Biotrophic Fungi.” The Arabidopsis Book / American Society of Plant Biologists 6 (October): e0115.

60. Michel, Elena J. S., Lalit Ponnala, and Klaas J. van Wijk. 2021. “Tissue-Type Specific Accumulation of the Plastoglobular Proteome, Transcriptional Networks, and Plastoglobular Functions.” Journal of Experimental Botany 72 (13): 4663–79.

61. Moellering, Eric R., Bagyalakshmi Muthan, and Christoph Benning. 2010. “Freezing Tolerance in Plants Requires Lipid Remodeling at the Outer Chloroplast Membrane.” Science 330 (6001): 226–28.

62. Murphy, Robert C. 2014. Tandem Mass Spectrometry of Lipids: Molecular Analysis of Complex Lipids. Royal Society of Chemistry.

63. Okazaki, Yozo, and Kazuki Saito. 2014. “Roles of Lipids as Signaling Molecules and Mitigators during Stress Response in Plants.” The Plant Journal: For Cell and Molecular Biology 79(4): 584–96.

64. Pfleger, Brian F., Michael Gossing, and Jens Nielsen. 2015. “Metabolic Engineering Strategies for Microbial Synthesis of Oleochemicals.” Metabolic Engineering 29 (May): 1–11.

65. Przybyla-Toscano, Jonathan, Mélanie Roland, Frédéric Gaymard, Jérémy Couturier, and Nicolas Rouhier. 2018. “Roles and Maturation of Iron-Sulfur Proteins in Plastids.” Journal of Biological Inorganic Chemistry: JBIC: A Publication of the Society of Biological Inorganic Chemistry 23 (4): 545–66.

66. Regmi, Anushobha, Jay Shockey, Hari Kiran Kotapati, and Philip D. Bates. 2020. “Oil-Producing Metabolons Containing DGAT1 Use Separate Substrate Pools from Those Containing DGAT2 or PDAT.” Plant Physiology 184 (2): 720–37.

67. Reuber, T. L., J. M. Plotnikova, J. Dewdney, E. E. Rogers, W. Wood, and F. M. Ausubel. 1998. “Correlation of Defense Gene Induction Defects with Powdery Mildew Susceptibility in Arabidopsis Enhanced Disease Susceptibility Mutants.” The Plant Journal: For Cell and Molecular Biology 16 (4): 473–85.

68. Saha, Saikat, Balaji Enugutti, Sona Rajakumari, and Ram Rajasekharan. 2006. “Cytosolic Triacylglycerol Biosynthetic Pathway in Oilseeds. Molecular Cloning and Expression of Peanut Cytosolic Diacylglycerol Acyltransferase.” Plant Physiology 141 (4): 1533–43.

69. Shimada, Takashi L., Makoto Hayashi, and Ikuko Hara-Nishimura. 2017. “Membrane Dynamics and Multiple Functions of Oil Bodies in Seeds and Leaves.” Plant Physiology 176 (1): 199– 207.

70. Shimada, Takashi L., Yoshitaka Takano, Tomoo Shimada, Masayuki Fujiwara, Yoichiro Fukao, Masashi Mori, Yozo Okazaki, et al. 2014. “Leaf Oil Body Functions as a Subcellular Factory for the Production of a Phytoalexin in Arabidopsis.” Plant Physiology 164 (1): 105– 18.

71. Shiva, Sunitha, Thilani Samarakoon, Kaleb A. Lowe, Charles Roach, Hieu Sy Vu, Madeline Colter, Hollie Porras, et al. 2020. “Leaf Lipid Alterations in Response to Heat Stress of Arabidopsis Thaliana.” Plants 9 (7). 10.3390/plants9070845.

72. Shiva, Sunitha, Hieu Sy Vu, Mary R. Roth, Zhenguo Zhou, Shantan Reddy Marepally, Daya Sagar Nune, Gerald H. Lushington, Mahesh Visvanathan, and Ruth Welti. 2013. “Lipidomic Analysis of Plant Membrane Lipids by Direct Infusion Tandem Mass Spectrometry.” Methods in Molecular Biology 1009: 79–91.

73. Smith, Matthew D., Donny D. Licatalosi, and John E. Thompson. 2000. “Co-Association of Cytochrome F Catabolites and Plastid-Lipid-Associated Protein with Chloroplast Lipid Particles1.” Plant Physiology 124 (1): 211–22.

74. Spanu, Pietro D. 2012. “The Genomics of Obligate (and Nonobligate) Biotrophs.” Annual Review of Phytopathology 50 (May): 91–109.

75. Springer, Armin, Chulhee Kang, Sachin Rustgi, Diter von Wettstein, Christiane Reinbothe, Stephan Pollmann, and Steffen Reinbothe. 2016. “Programmed Chloroplast Destruction during Leaf Senescence Involves 13-Lipoxygenase (13-LOX).” Proceedings of the National Academy of Sciences 113 (12): 3383–88.

76. Sutton, P. N., M. J. Henry, and J. L. Hall. 1999. “Glucose, and Not Sucrose, Is Transported from Wheat to Wheat Powdery Mildew.” Planta 208 (3): 426–30.

77. Swarbrick, Philip J., Paul Schulze-Lefert, and Julie D. Scholes. 2006. “Metabolic Consequences of Susceptibility and Resistance (race-Specific and Broad-Spectrum) in Barley Leaves Challenged with Powdery Mildew.” *Plant*, Cell & Environment 29 (6): 1061–76.

78. Tang, Dingzhong, Michael T. Simonich, and Roger W. Innes. 2007. “Mutations in LACS2, a Long-Chain Acyl-Coenzyme A Synthetase, Enhance Susceptibility to Avirulent Pseudomonas Syringae but Confer Resistance to Botrytis Cinerea in Arabidopsis.” Plant Physiology 144 (2): 1093–1103.

79. Tournayre, Jeremy, Matthieu Reichstadt, Laurent Parry, Pierre Fafournoux, and Celine Jousse. 2019. “‘Do My qPCR Calculation’, a Web Tool.” Bioinformation 15 (5): 369–72.

80. Tsitsigiannis, Dimitrios I., Terri M. Kowieski, Robert Zarnowski, and Nancy P. Keller. 2004. “Endogenous Lipogenic Regulators of Spore Balance in Aspergillus Nidulans.” Eukaryotic Cell 3 (6): 1398–1411.

81. Vallochi, Adriana Lima, Livia Teixeira, Karina da Silva Oliveira, Clarissa Menezes Maya-Monteiro, and Patricia T. Bozza. 2018. “Lipid Droplet, a Key Player in Host-Parasite Interactions.” Frontiers in Immunology 9 (May): 1022.

82. Vanhercke, Thomas, John M. Dyer, Robert T. Mullen, Aruna Kilaru, Md Mahbubur Rahman, James R. Petrie, Allan G. Green, Olga Yurchenko, and Surinder P. Singh. 2019. “Metabolic Engineering for Enhanced Oil in Biomass.” Progress in Lipid Research 74 (April): 103–29.

83. Vanhercke, Thomas, Anna El Tahchy, Pushkar Shrestha, Xue-Rong Zhou, Surinder P. Singh, and James R. Petrie. 2013. “Synergistic Effect of WRI1 and DGAT1 Coexpression on Triacylglycerol Biosynthesis in Plants.” FEBS Letters 587 (4): 364–69.

84. Vidi, Pierre-Alexandre, Marion Kanwischer, Sacha Baginsky, Jotham R. Austin, Gabor Csucs, Peter Dörmann, Felix Kessler, and Claire Bréhélin. 2006. “Tocopherol Cyclase (VTE1) Localization and Vitamin E Accumulation in Chloroplast Plastoglobule Lipoprotein Particles.” The Journal of Biological Chemistry 281 (16): 11225–34.

85. Wang, Liping, Wenyun Shen, Michael Kazachkov, Guanqun Chen, Qilin Chen, Anders S. Carlsson, Sten Stymne, Randall J. Weselake, and Jitao Zou. 2012. “Metabolic Interactions between the Lands Cycle and the Kennedy Pathway of Glycerolipid Synthesis in Arabidopsis Developing Seeds.” The Plant Cell 24 (11): 4652–69.

86. Wang, Wenming, Yingqiang Wen, Robert Berkey, and Shunyuan Xiao. 2009. “Specific Targeting of the Arabidopsis Resistance Protein RPW8.2 to the Interfacial Membrane Encasing the Fungal Haustorium Renders Broad-Spectrum Resistance to Powdery Mildew.” The Plant Cell 21 (9): 2898–2913.

87. Welti, Ruth, Weiqi Li, Maoyin Li, Yongming Sang, Homigol Biesiada, Han-E Zhou, C. B. Rajashekar, Todd D. Williams, and Xuemin Wang. 2002. “Profiling Membrane Lipids in Plant Stress Responses. Role of Phospholipase D Alpha in Freezing-Induced Lipid Changes in Arabidopsis.” The Journal of Biological Chemistry 277 (35): 31994–2.

88. Weng, Hua, Isabel Molina, Jay Shockey, and John Browse. 2010. “Organ Fusion and Defective Cuticle Function in a lacs1 lacs2 Double Mutant of Arabidopsis.” Planta 231 (5): 1089– 1100.

89. Weßling, Ralf, and Ralph Panstruga. 2012. “Rapid Quantification of Plant-Powdery Mildew Interactions by qPCR and Conidiospore Counts.” Plant Methods 8 (1): 35.

90. Wildermuth, Mary C. 2010. “Modulation of Host Nuclear Ploidy: A Common Plant Biotroph Mechanism.” Current Opinion in Plant Biology 13 (4): 449–58.

91. Wildermuth, Mary C., Michael A. Steinwand, Amanda G. McRae, Johan Jaenisch, and Divya Chandran. 2017. “Adapted Biotroph Manipulation of Plant Cell Ploidy.” Annual Review of Phytopathology 55 (August): 537–64.

92. Winter, Debbie, Ben Vinegar, Hardeep Nahal, Ron Ammar, Greg V. Wilson, and Nicholas J. Provart. 2007. “An ‘Electronic Fluorescent Pictograph’ Browser for Exploring and Analyzing Large-Scale Biological Data Sets.” PloS One 2 (8): e718.

93. Xu, Changcheng, Jilian Fan, and John Shanklin. 2020. “Metabolic and Functional Connections between Cytoplasmic and Chloroplast Triacylglycerol Storage.” Progress in Lipid Research 80 (November): 101069.

94. Xu, Changcheng, and John Shanklin. 2016. “Triacylglycerol Metabolism, Function, and Accumulation in Plant Vegetative Tissues.” Annual Review of Plant Biology 67 (April): 179–206.

95. Xue, Jinai, Huiling Gao, Yinghong Xue, Ruixiang Shi, Mengmeng Liu, Lijun Han, Yu Gao, et al. 2022. “Functional Characterization of Soybean Diacylglycerol Acyltransferase 3 in Yeast and Soybean.” Frontiers in Plant Science 13 (May): 854103.

96. Xu, Xiao-Yu, Hong-Kun Yang, Surinder P. Singh, Peter J. Sharp, and Qing Liu. 2018. “Genetic Manipulation of Non-Classic Oilseed Plants for Enhancement of Their Potential as a Biofactory for Triacylglycerol Production.” Proceedings of the Estonian Academy of Sciences: Engineering 4 (4): 523–33.

97. Yan, Bowei, Xiaoxuan Xu, Yingnan Gu, Ying Zhao, Xunchao Zhao, Lin He, Changjiang Zhao, Zuotong Li, and Jingyu Xu. 2018. “Genome-Wide Characterization and Expression Profiling of Diacylglycerol Acyltransferase Genes from Maize.” Genome / National Research Council Canada = Genome / Conseil National de Recherches Canada 61 (10): 735–43.

98. Yao, Hong-Yan, Yao-Qi Lu, Xiao-Li Yang, Xiao-Qing Wang, Zhipu Luo, De-Li Lin, Jia-Wei Wu, and Hong-Wei Xue. 2023. “Arabidopsis Sec14 Proteins (SFH5 and SFH7) Mediate Interorganelle Transport of Phosphatidic Acid and Regulate Chloroplast Development.” Proceedings of the National Academy of Sciences of the United States of America 120 (6): e2221637120.

99. Yin, Xiangzhen, Xupeng Guo, Lizong Hu, Shuangshuang Li, Yuhong Chen, Jingqiao Wang, Richard R-C Wang, Chengming Fan, and Zanmin Hu. 2022. “Genome-Wide Characterization of DGATs and Their Expression Diversity Analysis in Response to Abiotic Stresses in Brassica Napus.” Plants 11 (9). 10.3390/plants11091156.

100. Ytterberg, A. Jimmy, Jean-Benoit Peltier, and Klaas J. van Wijk. 2006. “Protein Profiling of Plastoglobules in Chloroplasts and Chromoplasts. A Surprising Site for Differential Accumulation of Metabolic Enzymes.” Plant Physiology 140 (3): 984–97.

101. Zhang, Meng, Jilian Fan, David C. Taylor, and John B. Ohlrogge. 2009. “DGAT1 and PDAT1 Acyltransferases Have Overlapping Functions in Arabidopsis Triacylglycerol Biosynthesis and Are Essential for Normal Pollen and Seed Development.” The Plant Cell 21 (12): 3885– 3901.

102. Zhao, Lifang, Vesna Katavic, Fengling Li, George W. Haughn, and Ljerka Kunst. 2010. “Insertional Mutant Analysis Reveals That Long-Chain Acyl-CoA Synthetase 1 (LACS1), but Not LACS8, Functionally Overlaps with LACS9 in Arabidopsis Seed Oil Biosynthesis.” The Plant Journal: For Cell and Molecular Biology 64 (6): 1048–58.

103. Zhou, Xue-Rong, Sajina Bhandari, Brandon S. Johnson, Hari Kiran Kotapati, Doug K. Allen, Thomas Vanhercke, and Philip D. Bates. 2019. “Reorganization of Acyl Flux through the Lipid Metabolic Network in Oil-Accumulating Tobacco Leaves1 [OPEN].” Plant Physiology 182 (2): 739–55.

104. Zhou, Xue-Rong, Pushkar Shrestha, Fang Yin, James R. Petrie, and Surinder P. Singh. 2013. “AtDGAT2 Is a Functional Acyl-CoA:diacylglycerol Acyltransferase and Displays Different Acyl-CoA Substrate Preferences than AtDGAT1.” FEBS Letters 587 (15): 2371– 76.

105. Zou, J., Y. Wei, C. Jako, A. Kumar, G. Selvaraj, and D. C. Taylor. 1999. “The Arabidopsis Thaliana TAG1 Mutant Has a Mutation in a Diacylglycerol Acyltransferase Gene.” The Plant Journal: For Cell and Molecular Biology 19 (6): 645–53.

106. Shiva, S., Vu, H. S., Roth, M. R., Zhou, Z., Marepally, S. R., Nune, D. S., Lushington, G. H., Visvanathan, M., & Welti, R. 2013). “Lipidomic analysis of plant membrane lipids by direct infusion tandem mass spectrometry.” *Methods in molecular biology (Clifton*, N.J.), 1009, 79–91. 10.1007/978-1-62703-401-2_9

